# Detecting brain network communities: considering the role of information flow and its different temporal scales

**DOI:** 10.1101/743732

**Authors:** Lazaro M. Sanchez-Rodriguez, Yasser Iturria-Medina, Pauline Mouches, Roberto C. Sotero

## Abstract

The identification of community structure in graphs continues to attract great interest in several fields. Network neuroscience is particularly concerned with this problem considering the key roles communities play in brain processes and functionality. Most methods used for community detection in brain graphs are based on the maximization of a parameter-dependent modularity function that often obscures the physical meaning and hierarchical organization of the partitions of network nodes. In this work, we present a new method able to detect communities at different scales in a natural, unrestricted way. First, to obtain an estimation of the information flow in the network we release random walkers to freely move over it. The activity of the walkers is separated into oscillatory modes by using empirical mode decomposition. After grouping nodes by their co-occurrence at each time scale, *k*-modes clustering returns the desired partitions. Our algorithm was first tested on benchmark graphs with favorable performance. Next, it was applied to real and simulated anatomical and/or functional connectomes in the macaque and human brains. We found a clear hierarchical repertoire of community structures in both the anatomical and the functional networks. The observed partitions range from the evident division in two hemispheres –in which all processes are managed globally– to specialized communities seemingly shaped by physical proximity and shared function. Our results stimulate the research of hierarchical community organization in terms of temporal scales of information flow in the brain network.

**Highlights:**

- Oscillatory modes of networks’ signals carry information on architectural rules.
- Meaningful partitions of the brain networks are found over different temporal scales.
- The multiscale organization of the brain responds to the function of its components.

## 1. Introduction

Network community detection constitutes a problem of current vital importance. Among all the nodes and interactions constituting a network, structures of subdivisions exist. In each of these communities (also referred to as groups or clusters), nodes have a greater probability of being locally connected than to nodes in other groups (Fortunato & Hric, 2016; Garcia, Ashourvan, Muldoon, Vettel, & Bassett, 2018). One example with several applications in the literature (Girvan & Newman, 2002; Porter, Mucha, Newman, & Warmbrand, 2005; Traud, Kelsic, Mucha, & Porter, 2011) is the tight-knit of a person’s friendships and the exchanges they have with other groups of friends. The identification of community structures provides insights into organizational principles, not only in terms of isolation of the clusters per se but also for the collective dynamical spreading of processes over the network (Fortunato & Hric, 2016).

In the brain, neural units connect to one another over different spatio-temporal scales in intriguing and fascinating ways (Christopher J Honey, Kotter, Breakspear, & Sporns, 2007; Moradi, Dousty, & Sotero, 2019; Sotero & Trujillo-Barreto, 2008; Valdes-Sosa et al., 2009). The modularity of such a system is believed to critically impact the phenomena of segregation (processes occurring in groups of heavily interconnected brain units) and integration (the combination of information exclusive to specialized brain regions) (Rubinov & Sporns, 2010; Sporns, 2013). Other advantages of a community structure relate to adaptability, robustness to failure and the reduction of wiring costs –see (Garcia et al., 2018) and (Betzel et al., 2017) and the references therein. Additionally, grouping exists across different levels (a hierarchy) for supporting rapid responses to changes (Garcia et al., 2018). As an illustration, consider the large community of neural conglomerates in one cerebral hemisphere. This can be broken into smaller communities according to the functional role of their members (Thomas Yeo et al., 2011). An important initial step for the study of brain structure, however, is the definition of the nodes and edges in the graph and the scale to be considered. This selection relies on the data available, which depends on the imaging modality used to record it. For example, anatomical associations can be examined through diffusion-weighted magnetic resonance imaging data (DWMRI) and functional neuroimaging or electrophysiological methods, e.g., fMRI and electroencephalogram provide insights into the dynamic interactions between brain regions (Y. Iturria-Medina et al., 2007; Valdes-Sosa et al., 2009).

Regardless of the network data, the bulk of community studies in the brain use variants of Newman’s modularity function (Newman, 2006) and its maximization through Louvain-like algorithms (Blondel, Guillaume, Lambiotte, & Lefebvre, 2008) for the detection of clusters of regions (Sporns, 2013). The partitions obtained through these methods maximize intra-community edge weights relative to a specific random network null model (Bassett et al., 2013; Garcia et al., 2018). Overall, these algorithms are problematic in that the output structure depends on the chosen null model and on a resolution parameter, *γ*, as well. Exploration of the resolution parameter space yields several structures that occasionally present hierarchy (Bassett, Khambhati, & Grafton, 2017). How could one set *γ* so that a meaningful set of communities –and not any partition– is revealed? In many instances, researchers exclusively report the partition obtained for *γ* = 1 (Fukushima et al., 2018). Nevertheless, it is known that high-modularity partitions can be found for *γ* = 1 in random, unstructured graphs, where no community structure should be detected (Fortunato & Hric, 2016). Recently, a useful heuristic has been introduced to retain the so-called graph’s most salient partition (Bassett et al., 2013; Garcia et al., 2018). In brief, a grid search is performed on the resolution parameter to find the value that generates the set of partitions with the greatest similarity. However, modularity maximization tends to split large communities into smaller pieces, which is a consequence of the choice of the null model. This effect is not solved by multi-resolution approaches (Fortunato & Hric, 2016). These techniques have also been adapted to generate hierarchical output structures (Ashourvan, Telesford, Verstynen, Vettel, & Bassett, 2019; Jeub, Sporns, & Fortunato, 2018) though the limitations with regard to the choice of null models and resolution parameters persist.

Other algorithms exist with somewhat fewer applications in brain research (Fortunato & Hric, 2016; Gates, Henry, Steinley, & Fair, 2016). Given the connectivity characteristics of communities, the utilization of random walkers for their identification is fairly straight-forward. Walkers tend to stay trapped in a cluster before transitioning to a different group (Fortunato & Hric, 2016). Walktrap (Pons & Latapy, 2005) and Infomap (M. Rosvall & Bergstrom, 2008) are examples of detection methods that employ random walk dynamics. The former is a costly, parameter-dependent method that exploits the probability of transition between two nodes in a certain number of steps as a measure of vertex similarity to group nodes. In the latter, a codeword is assigned to each vertex the walker encounters. Infomap considers networks with community structure to be analogous to geographical maps: unique codewords (street names) are only necessary to identify nodes (streets) in one specific community (city). Although Infomap has proven effective in artificial benchmark graphs and large datasets, it has performed more poorly in classical real networks traditionally utilized for testing algorithms (Gates et al., 2016; Hric, Darst, & Fortunato, 2014), e.g. the Zachary karate club (Girvan & Newman, 2002; Zachary, 1977). Those networks, for which ground-truth partitions are known, resemble some commonly analyzed brain graphs in that they have relatively small size and present various types of adjacency matrices, e.g., sparse, like those obtained from DWMRI or dense, from fMRI (Gates et al., 2016).

The description in terms of dynamical flows, as utilized in Walktrap and Infomap despite the above-mentioned limitations, is one that appeals to neuroscientists. In the first place, structural features like node degrees, the number of edges of the brain graph, etc., condition the dynamics of network processes (Fortunato & Hric, 2016). Secondly, the transmission of information in the brain is, obviously, a dynamical process (Sotero, Sanchez-Rodriguez, Dousty, Iturria-Medina, & Sanchez-Bornot, 2019), brain connectivity being adaptive and function-sensitive within the context of structural constraints (Friston, 2011). For these reasons, we believe that the analysis of community structure and the identification of hierarchical architectures in brain networks can benefit from considering the dynamical aspects of its information flow. Thus, in this paper, we present a novel approach to community detection specifically designed for brain graphs, although not limited to them.

We build on the methodology introduced by Sotero et al. in a recent work (Sotero et al., 2019). These authors studied information flow in brain networks by using the fraction of walkers that a given one finds at each node as the variable describing the evolution of the walker over the network. This function of time, taken as the network’s signal, can be decomposed into its constituent frequencies by using empirical mode decomposition (EMD) (N. E. Huang et al., 1996). Each of these oscillatory modes then associates with the notion of a temporal scale. Here, we incorporate a final step for performing data partitioning through *k*-modes clustering (Z. Huang, 1998). The arrangement of the nodes visits recorded throughout the walkers’ flows at the different temporal scales allows the unveiling of a hierarchical organization. Intuitively, a walker would spend considerable time in large communities, which is seen in slow oscillatory modes. Analogously, fast modes could reflect the motion over smaller clusters. Initially, we test the algorithm on benchmarks and real networks with known community structures such as Girvan-Newman (Girvan & Newman, 2002), Lancichinetti-Fortunato-Radicchi (Lancichinetti, Fortunato, & Radicchi, 2008) and the Zachary karate club (Zachary, 1977). We then proceed to extract communities existing in macaque (Christopher J Honey et al., 2007) and human anatomical connectivity matrices (Yasser Iturria-Medina, Sotero, Toussaint, & Evans, 2014), as well as *in-silico* (Cabral, Hugues, Sporns, & Deco, 2011) and *in-vivo* (Smith et al., 2015) functional connectivity graphs. Meaningful patterns of communities obtained here support the reliability of our method.

## 2. Materials and methods

### 2.1 The network’s signal

Let us imagine a network of *n* nodes, possibly presenting community structure, in which a random walker is set free. The walker moves over the edges available to it. In general, the probability of transitioning from node *j* towards node *i* in the next time step is given by (Zhang, Shan, & Chen, 2013):

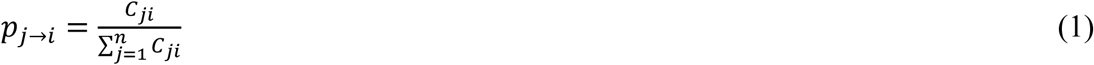

where *C*_*ji*_ is the weight of the connection from node *j* to node *i*. In other words, the process is described by Markov dynamics of first-order (the transition probabilities depend on the previous state) (Delvenne, Yaliraki, & Barahon, 2010). The walker tends to visit the nodes in a community before a route takes it to an outsider, a member of a different community –see (Fortunato & Hric, 2016) and the references therein. This is because of the predominantly local connectivity pattern of communities and the low number of outbound routes (Delvenne et al., 2010; Fortunato & Hric, 2016; M. Rosvall & Bergstrom, 2008; Martin Rosvall, Esquivel, Lancichinetti, West, & Lambiotte, 2014; Zhou, 2003). Many studies have extensively explored the particularities of community structure in terms of diffusion processes (Delvenne et al., 2010; M. Rosvall & Bergstrom, 2008; Martin Rosvall et al., 2014; Zhou, 2003). Specifically, (Delvenne et al., 2010) characterized the stability of partitions across time scales: “natural communities at a given time scale correspond to […] sets of states from which scape is unlikely within the given time scale”. Over short (fast) time scales, small clusters persist, whereas larger clusters arise as time evolves.

Now, suppose that *W*(*W* ≫ *n*) walkers simultaneously move over the same network. Each time one walker appears in a node, it finds fellow walkers, while others visit different nodes. Let us compute, for each walker, the fraction of the total number of other walkers that it encounters at each time step. After *T* time steps, there exist *W* time series reflecting different realizations of the flow of information in the network (Sotero et al., 2019). Those series incorporate information on the structure of the network (e.g., the number of walkers at a hub is expected to be persistently high), and the paths therein existing (i.e., the random walk itself). Finally, for generalization purposes –as the ratio of walkers would depend on the size of the network– we standardize such time series. Fig. 1a shows an exemplary signal corresponding to one of the walkers flowing over one of the networks considered in this paper. The horizontal axis is two-fold, showing both the temporal iteration (lower) and the indexes of the nodes the selected walker visits at each time step. Given the size of the graphs in this study, i.e., brain networks with *n*∼10^2^, we fix *W* = 1000.

**Fig. 1.**
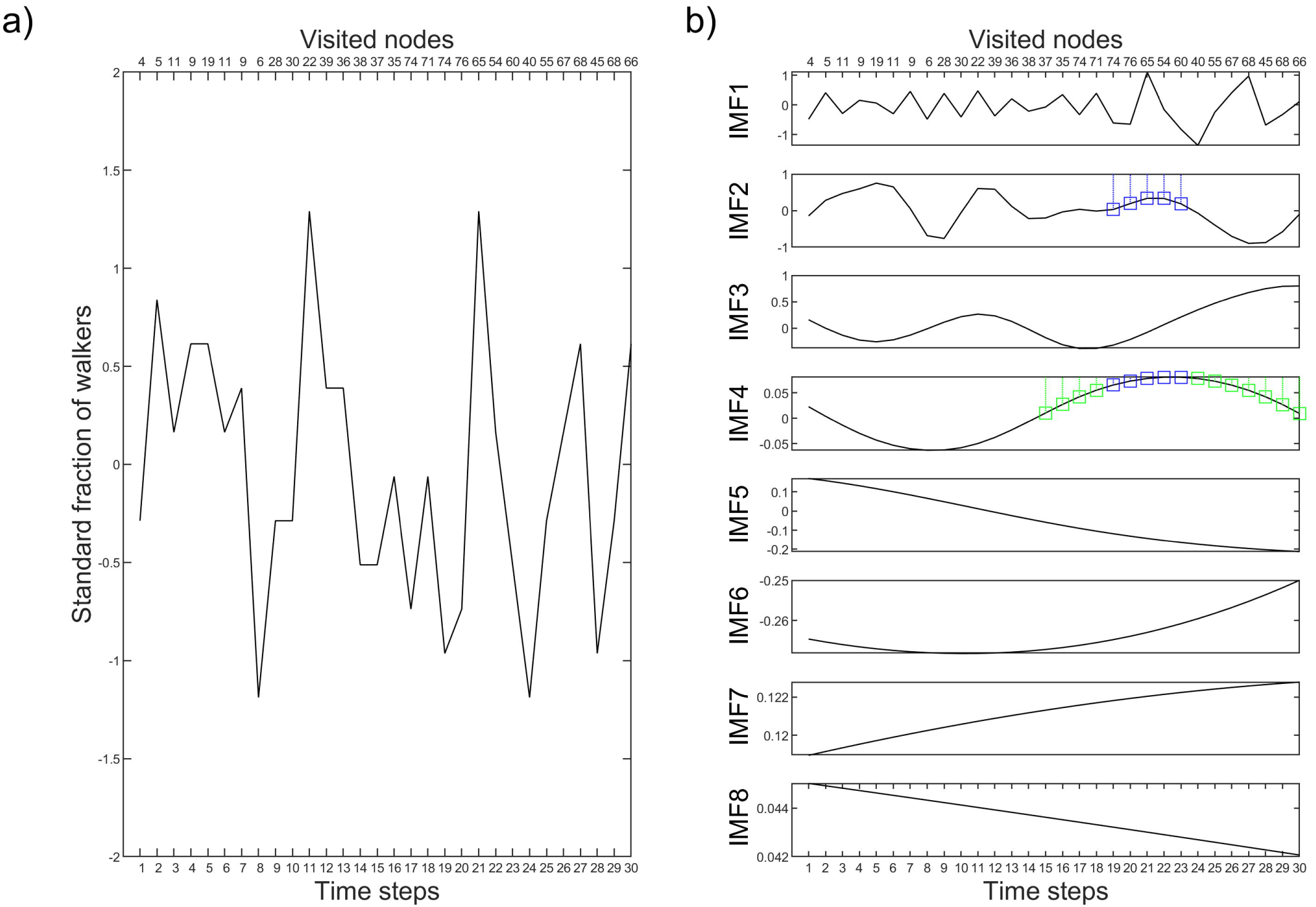
A typical network’s signal. a) Standard fraction of other random walkers one walker finds while flowing in a network. The lower horizontal axis shows the temporal progression of the walker. The upper axis shows the succession of nodes it visits. b) Empirical mode decomposition of the signal in (a). The activity of five nodes that are found between two zero-crossings of IMF_2_ is highlighted in blue. Together with those, other nodes appear between zero-crossings of IMF_4_ (in green). Only the first 30 time steps of *T* = 5000 are shown for visualization purposes.

The representation of the network we come up with can be decomposed to obtain oscillatory modes at different time scales. The temporal scales of random walks processes in complex networks depend on the network structure (Sotero et al., 2019). In other words, the network structure can be seen through different dynamical levels ranging from slow time scales in which walkers practically travel through the entire network to faster scales consisting, for instance, of the abrupt transitions from one node to the next. Empirical mode decomposition (EMD) (N. E. Huang et al., 1996) solves the problem of finding a nearly orthogonal basis for any complicated, nonlinear and non-stationary process without the need for any predefined model. The components into which the signal is broken down representing temporal scales are conventionally called intrinsic mode functions (IMFs). Each of these satisfy that: 1) the number of zero-crossings and extrema of the function are either equal or differ by one, and 2) the mean of its upper and lower envelopes is zero. Components are different as to conveyance of information (Sotero, 2016). For notation purposes, IMF_1_ denotes the fastest mode (highest frequency). The rest are named accordingly.

In our interpretation oriented to community detection, the IMFs of the signal corresponding to the fraction of walkers reflect hierarchical organization of the network. For example, fast temporal scales of the fraction of walkers could associate with partitions composed of small groups of nodes. In slower modes, the walker may get to visit all the nodes existing in larger communities. To give an example, as shown in Fig. 1b, a walker may quickly transition over nodes 74, 76, 65, 54 and 60 (seen with IMF_2_) or more slowly appear at those but also at others (IMF_4_). The five elements mentioned above may represent a community; those and the ones identified by the green color, may constitute a broader community.

### 2.2 Finding nodes that cluster together

The following step consists of exploiting the features of the IMFs and grouping nodes together. For each oscillatory mode and walker, we take chunks of data consisting of the network nodes seen between zero crossings. In our previous example, 74, 76, 65, 54 and 60 would be one of such data chunks for the fast IMF_2_ (Fig. 1b). Other sets of nodes will appear in different portions. One may think of our selection of the chunks in terms of the oscillations of a spring-mass system. There, points to the right/left of the zero-reference cluster together (the spring is stretched/compressed). Each time the signal for the displacement of the mass passes through the equilibrium position it is also switching from one ‘community’ to the other. An animation of the oscillations of the spring-mass system is shown in the Supplementary Video.

#### Supplementary video goes around here

**Supp. video**. The oscillations of a spring-mass system. Points to the right/left of the zero-reference cluster together (the spring is stretched/compressed). When the signal for the displacement of the mass (right panel) passes through the equilibrium position it is also switching from ‘the community of positive coordinates’ to ‘the community of negative coordinates’ and vice versa.

The zero crossings-analogy bases on the interpretation and symmetry properties of the IMFs. In practice, nodes outside a certain true community *C*_*q*_, may occasionally pertain to a chunk of otherwise genuine members of *C*_*q*_, given the existence of edges to that community. Likewise, all nodes belonging to a community do not necessarily have to appear together between two contiguous zero crossings. The problem is how to identify authentic node clusters over the effects of noise with the information available from the IMFs. To address this issue, we turn to unsupervised learning, particularly to clustering. In clustering analysis, the goal is to group objects based on the information available –features describing the data (Assent, 2012; Ronan, Qi, & Naegle, 2016; Steinbach, Ertöz, & Kumar, 2004). The more similar items in a group are and the more different to those in other groups, the better the clustering (Steinbach et al., 2004).

To capture the natural structure of the data, we proceed to cluster network nodes –or objects, in conventional clustering jargon– given their co-appearance between zero-crossings of the IMFs –the features. Features are binary vectors encoding the positions of the nodes appearing together in the between zero-crossings chunks. In other words, we think of each of the IMFs as independent ideal oscillators whose zero-crossings separate various communities of nodes, the clustering algorithm being the tool that resolves the natural shortcomings of this abstraction. A complete representation of the IMFs corresponding to each walker (Fig. 1) in terms of binary variables is thus obtained (Fig. 2a). To ensure a proper sampling of all nodes and their co-appearances, we select the features corresponding to a high number of walkers (200 out of the 1000 simulated). This is a random selection, in the same way that subsets of variables are sometimes chosen when clustering data with multiple independent signals (Jiliang Tang, Salem Alelyani, & Huan Liu, 2014; Ronan et al., 2016).

**Fig. 2.**
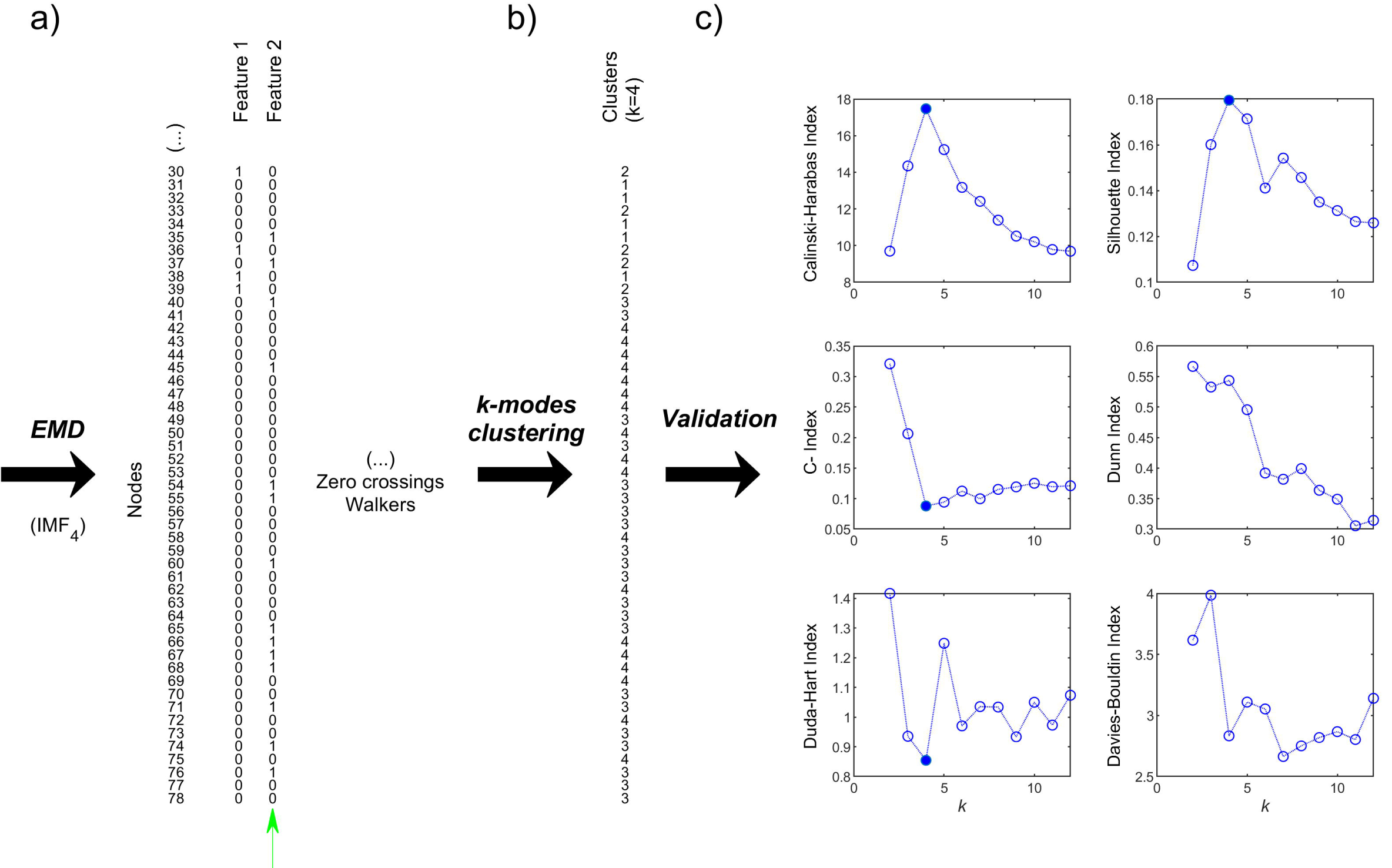
Clustering network nodes. a) -For each IMF, ‘features’ are constructed so that nodes appearing together between zero-crossings of the IMF are assigned a logical 1. For example, Feature 2 here corresponds to the highlighted nodes for IMF_4_ of Fig. 1b. Such description is extended through all zero-crossings of one IMF and to other walkers to guarantee a proper sampling of the network nodes and their close acquaintances in the temporal scale. b) The obtained data matrix feeds a *k*-modes clustering algorithm. For one instance of the data and *k* = 4 clusters, the algorithm returns the solution shown (e.g. nodes 76, 77 and 78 are in “community 3”). c) After exploring the *k*-space, all the solutions are considered according to several validation indexes. In the example, the Calinski-Harabasz index and the silhouette width, both presenting an absolute maximum, together with C-index and the Duda-Hart measure (having absolute minimums), suggest the existence of 4 clusters in the data. This is sufficient to come to a conclusion although the resting indexes (Dunn and Davies-Bouldin) present relative maximum and minimum at *k* = 4, respectively, which according to their definition may as well indicate the presence of network organization in 4 communities. Values of *k* for which *k*-modes yields singleton communities are not shown (*k* = [13, 20], *k* ∈ ℕ).

Here, we employ the so-called *k*-modes clustering algorithm (Z. Huang, 1998). *K*-modes is an extension of the popular *k*-means method (Macqueen, 1967) for categorical variables, binary features being a particular case of those. A simple matching dissimilarity is used as notion of distance in *k*-modes. Two objects, ***x*** and ***y*** are far from each other by a quantity that equals the number of mismatching features (of *M* features), namely:

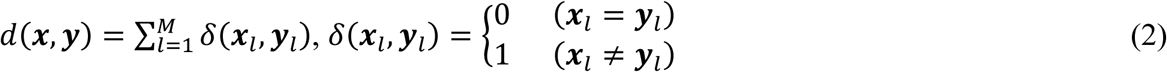

The algorithm minimizes a cost function after a vector (the mode^1^) for each of the *k* clusters has been selected and objects grouped around it such that their dissimilarity is minimal. Alike *k*-means, Huang’s *k*-modes yields locally optimal solutions depending on the starting conditions. Thus, one necessary first step is running the algorithm several times to select the solution with the lowest overall cost. The number of initial conditions for the clustering algorithm is empirically set to 50 in this work, based on the consistency of the solutions obtained. With all these considerations *k*-modes is run (Fig. 2b). The implementation of *k*-modes we used is available from https://github.com/nicodv/kmodes.

### 2.3 Accepting/rejecting hierarchical partitions

Clustering algorithms generally find clusters even in the absence of underlying structure, highlighting the necessity of validating solutions (Ronan et al., 2016). In *k*-modes, objects are allocated in *k* clusters exactly. However, unless the user has prior knowledge on the distribution of the data –which rarely occurs– *k* is a parameter to be determined. Several metrics are intended to elucidate the correct number of clusters in the data from running the algorithm over a range of *k*. These metrics use measures of separation, compactness, or both. Various studies, most famously the one by Milligan and Cooper (Milligan & Cooper, 1985), have looked at the performance of indexes for assessing the results over numerical data and Euclidean distances. To the best of our knowledge, such measures have not been transformed to account for categorical data (binary, in particular) and matching dissimilarity, as recommended by Huang in his seminal paper (Z. Huang, 1998). Therefore, in Appendix B, we briefly describe six measures with satisfactory performance for recovering true cluster structure, i.e.: the Calinski-Harabasz index (Anderson, 2001; Calinski & Harabasz, 1974), the C-index (Milligan & Cooper, 1985), a modified Duda-Hart criterion (Duda, Hart, & Stork, 2001), silhouette width (Kaufman & Rousseeuw, 1990), one of the family of Dunn indexes (Bezdek & Pal, 1998; Dunn, 1973) and the Davies-Bouldin index (Davies & Bouldin, 1979; Dubes, 1987). Details on their utilization and customization to account for binary data, if applicable, are also included. Importantly, as the success ratio of an individual index in determining the true number of clusters is limited and may depend on the data (Dubes, 1987; Milligan & Cooper, 1985), here, we adopt the criterion of selecting the best number of clusters based on a majority rule (Charrad, Ghazzali, Boiteau, & Niknafs, 2014). Failure to establish a majority of the indexes indicating the same correct number of clusters may hint at the lack of a definitive community pattern. This is usually accompanied by at least one of the indexes rejecting the existence of community structure altogether (see Appendix C). Fig. 2c shows an example in which most of the validation measures indicate the existence of *k* = 4 clusters, to finalize the illustration of the basic pipeline of our method. For the networks considered in this paper, distributions of nodes in up to 20 clusters only were investigated.

The process illustrated in this figure should be performed for many combinations of walkers (to build consensus partitions) and for all the IMFs (to unveil hierarchical organization).

The exploration over values of *k* should be performed for each of the IMFs. The following intuitive rule of thumb is followed. First, take the slowest IMF in the decomposition of the signal corresponding to the fraction of walkers. After running the clustering algorithm, determine the number of communities existing. Move on to the next IMF and determine its corresponding partition and the number of clusters present. If this number is equal to any obtained beforehand, reject the previous partition and accept only the current one. Else, retain both partitions (with different numbers of clusters). Proceed with the analysis until all the IMFs have been considered. This way, a hierarchical organization is unveiled.

One last issue to consider is the stochasticity that is inherent to most community detection techniques (Bassett et al., 2013; Fortunato & Hric, 2016). In our method, this is expressed as sets of randomly chosen walkers which are considered to build clustering features. The application of the clustering algorithm over those subsets of features can yield slightly different partitions. To present unique partitions, we use consensus clustering (Strehl & Ghosh, 2003). One starts by building a consensus matrix, ***T***, that accounts for the co-occurrence of nodes in communities. Non-significant relationships between nodes are removed by thresholding the co-occurrence matrix. Such a threshold is set to the highest value of all co-occurrences in the association matrix resulting from random permutations of the original partitions (Bassett et al., 2013). Then, the algorithm is applied over ***T*** until all the partitions are identical (the ‘true-partition’). The results reported in this paper correspond to the consensus partition after applying *k*-modes clustering to 50 sets of independent features for each of the networks. Their similarity with known ground-truth communities is analyzed by using adjusted mutual information, AMI (N. Vinh, Epps, & Bailey, 2010) (see Appendix D).

### 2.4 Data description and processing

#### 2.4.1 Benchmarks

Artificially generated graphs and a real network with known group structure were used to assess the performance of our algorithm. We chose simple benchmark graphs with features alike the brain networks for which the application was intended, e.g. a similar number of nodes.

##### 2.4.1.1 Girvan-Newman benchmarks (GN)

These graphs are random with known community structure. A GN benchmark consists of 128 nodes and four communities, with 32 nodes each. The average expected degree of a node is 16 (Girvan & Newman, 2002). A fraction of those connections (*μ*) is made to vertices in other communities. As such fraction increases, algorithms usually struggle to pinpoint the underlying community structure. Here, we set *μ* = 0.1 (Lancichinetti & Fortunato, 2009) and applied our detection algorithm over both binary and weighted versions of the GN model. One limitation of GN is its inability to reproduce the scale-free property of real networks (heterogeneous node degree distributions, node degree and community sizes following a power-law distribution) (Lancichinetti et al., 2008).

##### 2.4.1.2 Lancichinetti-Fortunato-Radicchi benchmarks (LFR)

A more realistic benchmark, LFR does account for the heterogeneous and skewed distribution of the degree and community size. Both these parameters are chosen from power-law distributions. Networks are built by joining stubs at random (Fortunato & Hric, 2016; Lancichinetti et al., 2008). We kept the value of the mixing parameter at 0.1. The average degree in a network of *n* = 100 nodes was set to 13 and the upper extreme of the degree distribution to 27. Consequently, the randomly generated networks (binary and weighted) utilized here had 5 communities, with sizes [26, 23, 21, 20, 10].

The GN and LFR networks used in this work were generated by using code available from (https://www.santofortunato.net/resources, *LFR benchmark graphs*). All the code parameters were set to their default values except for the ones above-mentioned.

##### 2.4.1.3 Zachary’s karate club

The karate club network collects the interactions of 34 individuals over three years (Zachary, 1977). A conflict over the price of the karate lessons escalated and provoked the fission of the group as the supporters of the club’s instructor formed a new organization, separate from the original one that stayed with the president. Thus, Zachary’s data encompasses one of the few examples of nearly-definitive ground-truth communities, the two resulting groups (Hric et al., 2014). Many of the detection algorithms existing in the literature are tested on their ability to recuperate Zachary’s factions. Such a task is usually performed over a binary connectivity matrix for the members of the club (Girvan & Newman, 2002; Hric et al., 2014; Newman, 2004). In this study, instead, we used the weighted version provided by Zachary, in which the strength of an edge is given by the number of external contexts where interactions between two individuals were observed (see Appendix A, Fig. S1, for the adjacency matrix that fixes some inconsistencies in Zachary’s report). The weights were normalized to the [0,1] interval.

#### 2.4.2 Brain networks

We investigated the community structure of multiple brain networks, including macaque and human structural connectomes, simulated functional interactions based on Kuramoto oscillators (Cabral et al., 2011) and resting-state functional networks in the healthy human brain.

##### 2.4.2.1 Macaque visual and sensorimotor anatomical network

Cortico-cortical connections existing between large-scale areas of the macaque neocortex have been identified through anatomical tracing studies (Malcolm P Young, 1993). Among all the areas and pathways summarized in Young’s paper, only those lying in the cortical visual and somatosensory-motor systems are considered here (see Fig. S2). This connectivity matrix, with 46 nodes, is only slightly different than the one utilized in the network structure study by Honey et al. (Christopher J Honey et al., 2007), where visual areas were labeled following (Felleman & Van Essen, 1991). Several connections are reciprocal. However, in general, the network is directed and binary, with 1’s in a row indicating the efferent projections reported for the given area –see (Van Essen & Felleman, 1991; M P Young, 1993) for more details on the cortical areas.

##### 2.4.2.2 Human brain anatomical network

An average human brain anatomical network (Yasser Iturria-Medina et al., 2014) was also constructed and analyzed in this study. The original data is freely available by The Cognitive Axon (CoAx) Lab, in the Center for the Neural Basis of Cognition and Department of Psychology at Carnegie Mellon University (http://www.psy.cmu.edu/~coaxlab/data.html), who acquired and processed the data. Participants in the study included 60 subjects (29 males and 31 females; ages 18 to 45 years, mean 26 ± 6), recruited from the local Pittsburgh community and the Army Research Laboratory in Aberdeen Maryland. All subjects were neurologically healthy, with no history of either head trauma or neurological or psychiatric illness.

###### Ethics statement

The procedure was approved by the institutional review board at Carnegie Mellon University. Participants provided informed consent to participate in the study and consent to publish any research findings based on their provided data (Dunovan, Lynch, Molesworth, & Verstynen, 2015).

###### Image acquisition

Participants were scanned on a Siemens Verio 3T system in the Scientific Imaging & Brain Research (SIBR) Center at Carnegie Mellon University using a 32-channel head coil. Image collection was performed with the following parameters: 50 min, 257-direction DSI scan using a twice-refocused spin-echo EPI sequence and multiple q values (TR = 9916 ms, TE= 157 ms, voxel size = 2.4×2.4×2.4 mm, FoV = 231×231 mm, b-max = 5,000 s/mm^2^, 51 slices). Head movement was minimized during the scan.

###### Image processing

All images were processed using a q-space diffeomorphic reconstruction method (Yeh & Tseng, 2011) to register the voxel coordinates into MNI space (Evans, Kamber, Collins, & MacDonald, 1994). As a result of the processing across all 60 subjects, a final template image (CMU-60 DSI) was created by averaging the ODF maps. This template constitutes a detailed and unbiased representative map of the nervous fiber orientations in the young healthy brain.

Next, we estimated probabilistic axonal connectivity values between each brain voxel and the surface of each considered gray matter region (voxel-region connectivity) using a fully-automated fiber tractography algorithm (Y. Iturria-Medina et al., 2007) and the intravoxel fiber ODFs of the CMU-60 DSI Template. The tracking parameters were imposed as follows: a maximum of 500 mm trace length and a curvature threshold of ±90°. The anatomical regions were defined following the labeling procedure by Klein & Tourville (Klein & Tourville, 2012), from which 78 regions were considered –see (Y Iturria-Medina et al., 2016; Sanchez-Rodriguez et al., 2018) for more details. Based on the resulting voxel-region connectivity maps, the anatomical connection probability between any pair of regions *i* and *j* (0 ≤ *ACP*_*ji*_ ≤ 1, *ACP*_*ji*_ = *ACP*_*ij*_) was calculated as the maximum voxel region connectivity value between both regions. For any pair of regions *i* and *j*, the *ACP*_*ji*_ measure (Y. Iturria-Medina et al., 2007) reflects the degree of evidence supporting the existence of the hypothetical white matter connection, independently of the density/strength of this connection. A network backbone, containing the dominant connections in the regional connectivity map, was computed using a minimum-spanning-tree based algorithm (Rubinov & Sporns, 2010). It was the resulting minimum spanning tree the network that we used (Fig. 3a).

**Fig. 3.**
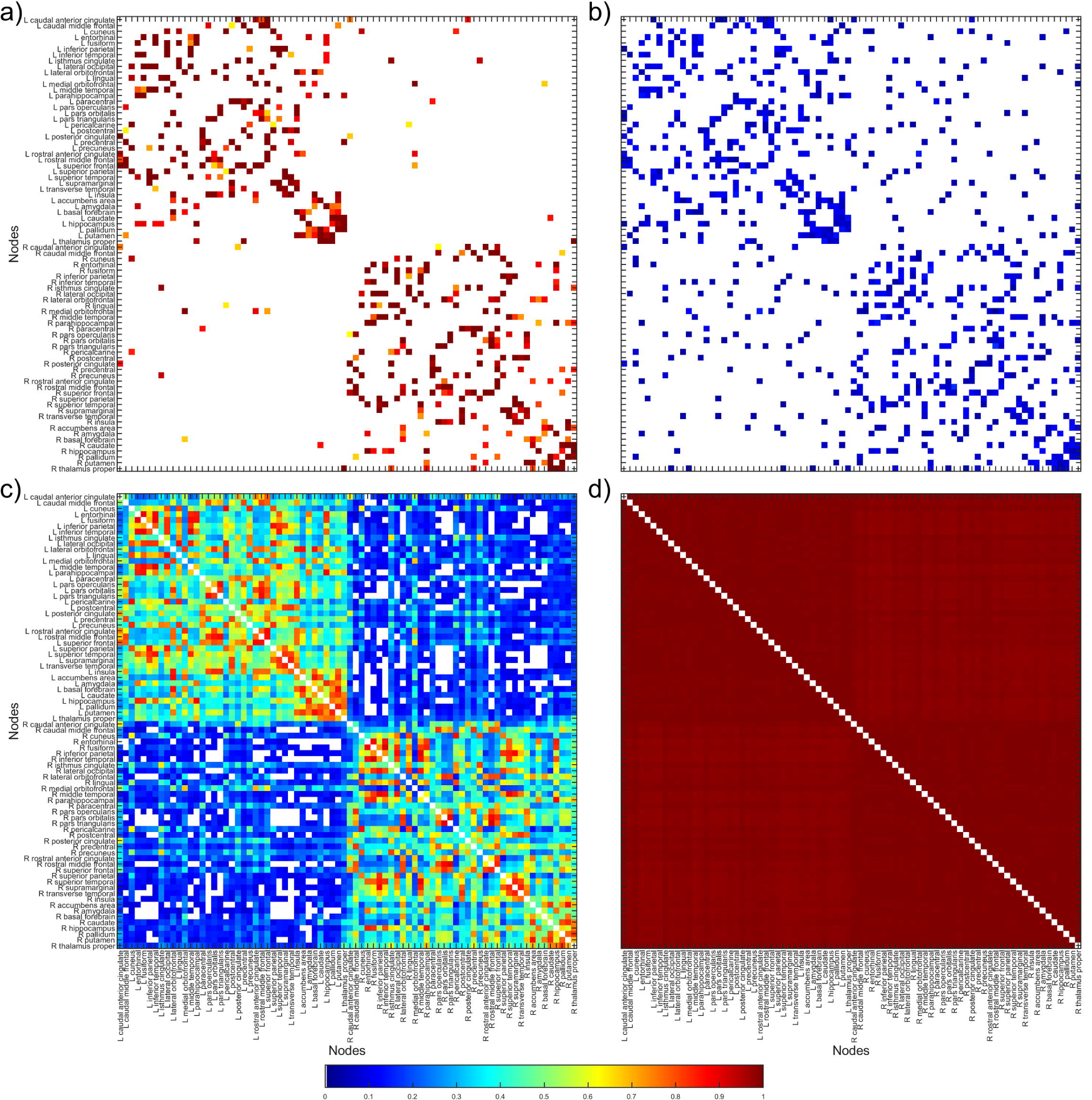
Human brain networks of the 78-regions anatomical parcellation. a) Anatomical connections between 78 brain areas. b) Functional connectivity obtained from superimposing Kuramoto oscillators to the matrix in (a). The global coupling strength is *κ* = 5. c) As in (b), with *κ* = 30. d) As in (b) and (c), with *к* = 150.

##### 2.4.2.3 Simulated human brain functional network

To construct a representation of functional interactions in the brain, simulations of the Kuramoto model (Kuramoto, 1975) were performed. The Kuramoto model is a classical dynamical system that describes the behavior of a set of coupled oscillators. For the sake of consistency and contrast, the anatomical parcellation described in the previous section was conserved, while the relative coupling between two nodes in the network of oscillators corresponded to the backbone-*ACP* measure between regions *j* and *i*. The evolution of the phase of the *i*-th oscillator, *θ*_*i*_, is given by (Daffertshofer & van Wijk, 2011):

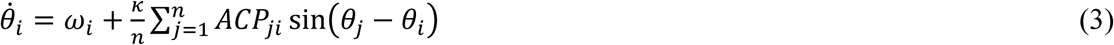

where *к* is a global coupling strength and *ω*_*i*_ is the intrinsic frequency of node *i*. In all our simulations, we drew the natural frequencies from a standard Gaussian distribution and the initial conditions from a uniform distribution in the interval [0, 2π). A total of 250 sets of natural frequencies and initial conditions were used. System (3) was numerically solved via a Euler scheme, with time step Δ*t* = 0.001*s* and *t*_*total*_ = 50*s*. The first simulated 10*s* were discarded in all occasions to reduce the effect of transients in the results.

Intuitively, the collective behavior of the system depends on the parameter *κ*. Stronger interactions (high *κ*) overcome the dispersion of the intrinsic frequencies yielding coherence in the network, whereas in the low−*κ* regime oscillators tend to remain asynchronous (Breakspear, Heitmann, & Daffertshofer, 2010; Daffertshofer & van Wijk, 2011). The degree of synchrony of the oscillators is quantified through the phase uniformity (K. V. Mardia, 1975):

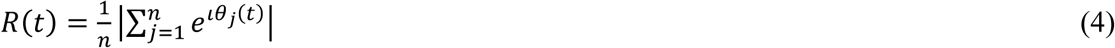

In our calculations, a grand-average phase uniformity value for each coupling strength, *κ*, was obtained by averaging *R*(*t*) in the considered time interval across all simulations with such *κ* (Fig. S3). Similarly, a so-called *κ*-dependent functional connectivity matrix was also calculated. To do so, we followed Cabral et al. (Cabral et al., 2011) and assumed an electrophysiological measure of the brain activity, such as the mean firing rate or the excitatory postsynaptic potential over the brain region, to be given as *y*_*i*_(*t*) = *y*_0_ sin(*θ*_*i*_(*t*)). Functional connectivity for a pair of nodes was then defined as the Pearson correlation between their *y*_*i*_(*t*) and *y*_*j*_(*t*) signals, for each simulation. The representative interaction matrix associated with *κ* was finally obtained after Fisher-transforming the pairwise correlation coefficients, averaging, performing one-sample *t*-tests (*α* = 0.05), correcting by false-discovery rate and applying the inverse transformation (Bordier, Nicolini, & Bifone, 2017). Functional connectivity matrices for *κ* = 5 (*R* ≈ 0.12), *κ* = 30 (*R* ≈ 0.47) and *κ* = 150 (*R* ≈ 0.98) are shown in Fig. 3b, Fig. 3c and Fig. 3d, respectively.

##### 2.4.2.4 Human brain resting-state fMRI network

Organization of the human brain in terms of functional connectivity was studied by using resting-state fMRI data and a parcellated connectome obtained by means of independent component analysis (ICA) (Mckeown et al., 1998), and available from the Human Connectome Project (HCP). The HCP500 “PTN” (Parcels, node Timeseries and Netmats) dataset used in this study (https://www.fmrib.ox.ac.uk/datasets/HCP-CCA/) was publicly released via the central HCP ConnectomeDB database in late 2014. Access to the HCP data is available online through https://db.humanconnectome.org/. We provide a short description of the dataset, following (Smith et al., 2015), and additional processing steps. Further details can be found elsewhere (Smith et al., 2013; Uğurbil et al., 2013; Van Essen et al., 2013).

###### Data

Resting-state fMRI data (rfMRI) was acquired from 461 healthy subjects (190 males and 271 females; ages 22 to 35 years) on a 3-T Siemens connectome-Skyra scanner. Each of the four 15-minutes runs of each subject had temporal resolution 0.73 s and spatial resolution 2-mm isotropic. T1-weighted and T2-weighted structural images of resolution 0.7-mm isotropic and B0 field mapping were also carried out (Smith et al., 2015).

###### Pre-processing and Group-ICA

Data were preprocessed according to (Smith et al., 2013) and had artifacts removed using ICA+FIX (Griffanti et al., 2014). The rfMRI runs were temporally demeaned and had variance normalization applied (Beckmann & Smith, 2004). Group-principal component analysis (PCA) output was generated by MIGP (MELODIC’s Incremental Group-PCA), comprising the top 4500 weighted spatial eigenvectors (Smith et al., 2015). The MIGP output was fed into group-ICA using FSL’s MELODIC tool (Beckmann & Smith, 2004), applying spatial-ICA at several different ICA dimensionalities, *n*. Spatial-ICA was applied in grayordinate space (surface and subcortical grey matter voxels) with volumetric MNI152 3D-space versions of these maps being available (see below).

###### Network matrices

For a given group-ICA decomposition, the set of ICA spatial maps was mapped onto each subject’s rfMRI time-series data to derive one representative time-series per ICA component. Effectively, each ICA component is considered as a network “node”. The node time-series were obtained by estimating the principal eigen-time-series within each ICA component (O’Reilly, Beckmann, Tomassini, Ramnani, & Johansen-Berg, 2010). Network matrices were obtained from the node time-series. Network modeling was performed through the FSLNets toolbox (https://fsl.fmrib.ox.ac.uk/fsl/fslwiki/FSLNets). For each subject and ICA dimensionality, the HCP provides a connectivity matrix where the strength of any connection is estimated using “full” normalized temporal correlation between every node time-series and every other (Smith et al., 2013).

###### Dimensionality and Group average connectivity matrix

A set of ICA maps can be considered a parcellation (Smith et al., 2015). The dimensionality determines the number of distinct ICA components; a low number typically means that the regions within the spatial component maps will be bigger. Each ICA map (node in the parcellation) is given as a set of non-overlapping voxels in MNI152 space in the file ‘melodic_IC_sum’ of the HCP500 “PTN” release. In this work, we picked *n* = 25 from all the dimensionalities considered by the HCP. This is i) to expeditiously illustrate the applicability of the methodology and ii) to facilitate the display and interpretability of the results through comparison with well-established resting-state sub-systems (Thomas Yeo et al., 2011). To assign each node to one of these sub-systems, we first co-registered and resampled the 17-sub-systems solution of (Thomas Yeo et al., 2011), available from https://surfer.nmr.mgh.harvard.edu/fswiki/CorticalParcellation_Yeo2011, to the space of volumetric ICA maps using *spm_reslice*.*m* in SPM12 (https://www.fil.ion.ucl.ac.uk/spm/). The greatest overlap (number of common voxels) was used to define a node’s assignment to a resting-state sub-system (Betzel, Gu, Medaglia, Pasqualetti, & Bassett, 2016; Sacchet et al., 2016). This correspondence is shown in Table S1. Nodes 17, 18 and 23 did not present any significant overlap with the 17 resting-state sub-systems and were eliminated from the analysis. Finally, a group connectivity matrix was obtained by averaging the Pearson correlation individual matrices as in section 2.4.2.3 (see *Methods, Simulated human brain functional network*) and is shown in Fig. S4.

#### 2.4.3 Data and code availability statement

The datasets and codes analyzed during the current study are available from public repositories, which have been referenced throughout the paper. A specific set of codes containing a demonstration on how to concatenate the method pipeline is offered at https://www.soterolab.com/software. All calculations but the detection of clusters were performed in MATLAB R2019a (The MathWorks Inc., Natick, MA, USA). Python 2.7 (Python Software Foundation, https://www.python.org/) was used for an implementation of the *k*-modes algorithm.

## 3 Results

### 3.1 Community detection in benchmark graphs

Table 1 shows the results of the application of our method in benchmark graphs (see *Methods, Benchmarks*). The AMI value (see *Appendix D*), appearing in the last column, illustrates the degree of similarity between the obtained partitions and the ground-truth community structure known for each of the graphs. For both instances of the GN model (binary and weighted), the right partition was found over a range of IMFs. In the case of the LFR benchmarks, our method unveiled the 5 communities planted at IMF_6_. Over slower IMFs than the ones reported in Table 1, the coarser organization of the networks was in some cases observed, e.g. one of the ground-truth communities stood alone and the rest merged. The analysis of faster IMFs did not return any community structure (see *Appendix B* and *Appendix C*). Finally, in the case of the karate club, the two known fractions in which it split were nearly obtained over IMF_3_.

**Table 1.**
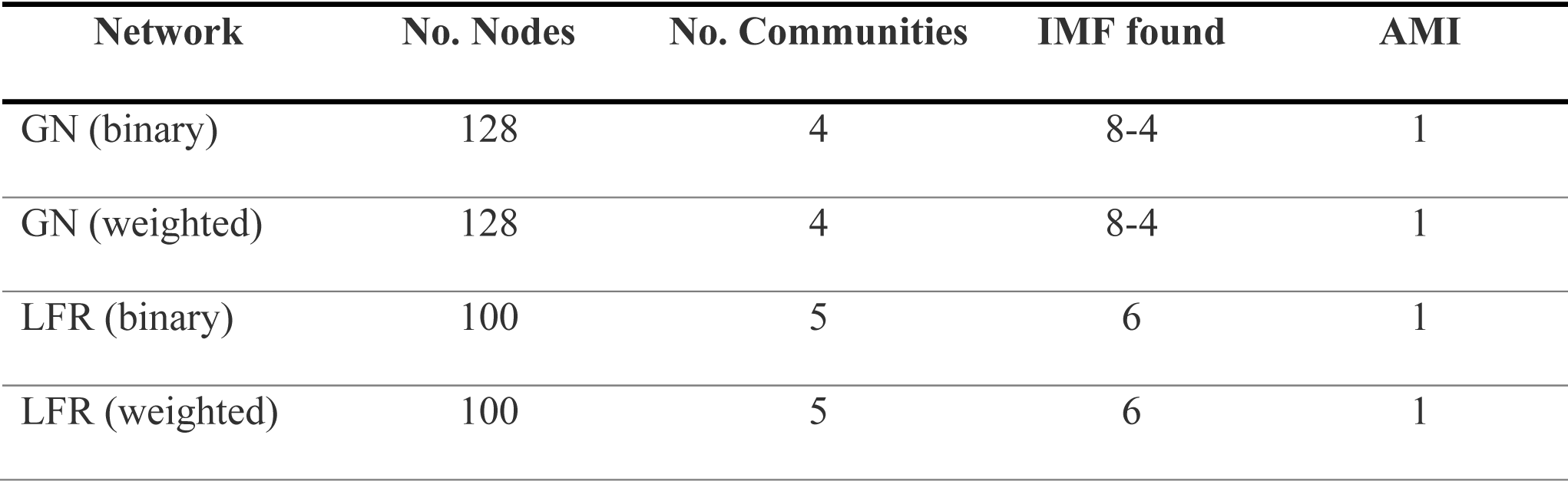
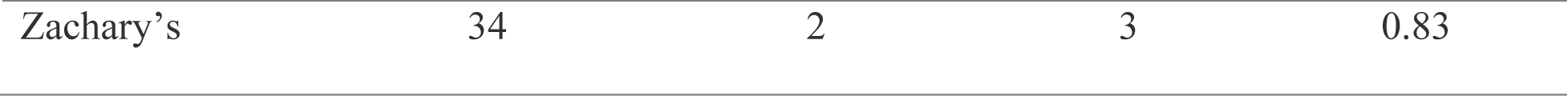
Characteristics of the benchmarks and results of the application of our detection method. For each network, the number of nodes and known communities existing are given. The IMFs over which our method finds the right number of communities appear in the fourth column. In the last column, the adjusted mutual information values quantifying the degree of similarity between the solutions returned by the algorithm and the ground-truth structures are shown.

The results for the network of Zachary’s karate club are further illustrated in Fig. 4. A schematic representation of the two-communities structure that was revealed appears in Fig. 4a. Fig. 4b contains information regarding the validation measures, showing the selection, by a majority rule, of two clusters in the data (see *Appendix B*). Likewise, other runs of the algorithm signaled the existence of two clusters. The left panel in Fig. 4c shows 50 of such partitions (one per row). In some of those, node 10 was assigned to the community we have called “1” (in blue). Thus, a consensus matrix (Fig. 4c, center panel) basically consists of binary values for the co-occurrences of all nodes in communities but those including node 10. Re-running the algorithm yielded 50 identical partitions (Fig. 4c, right). This partition (Fig. 4a) corresponds to the division reported by Zachary through observations of the karate club except for one member (node 9). This result is expected, according to the original paper and many others in which the karate club has been analyzed (Girvan & Newman, 2002; Hric et al., 2014), as the data apparently supports node 9’s membership to the wrong faction.

**Fig. 4.**
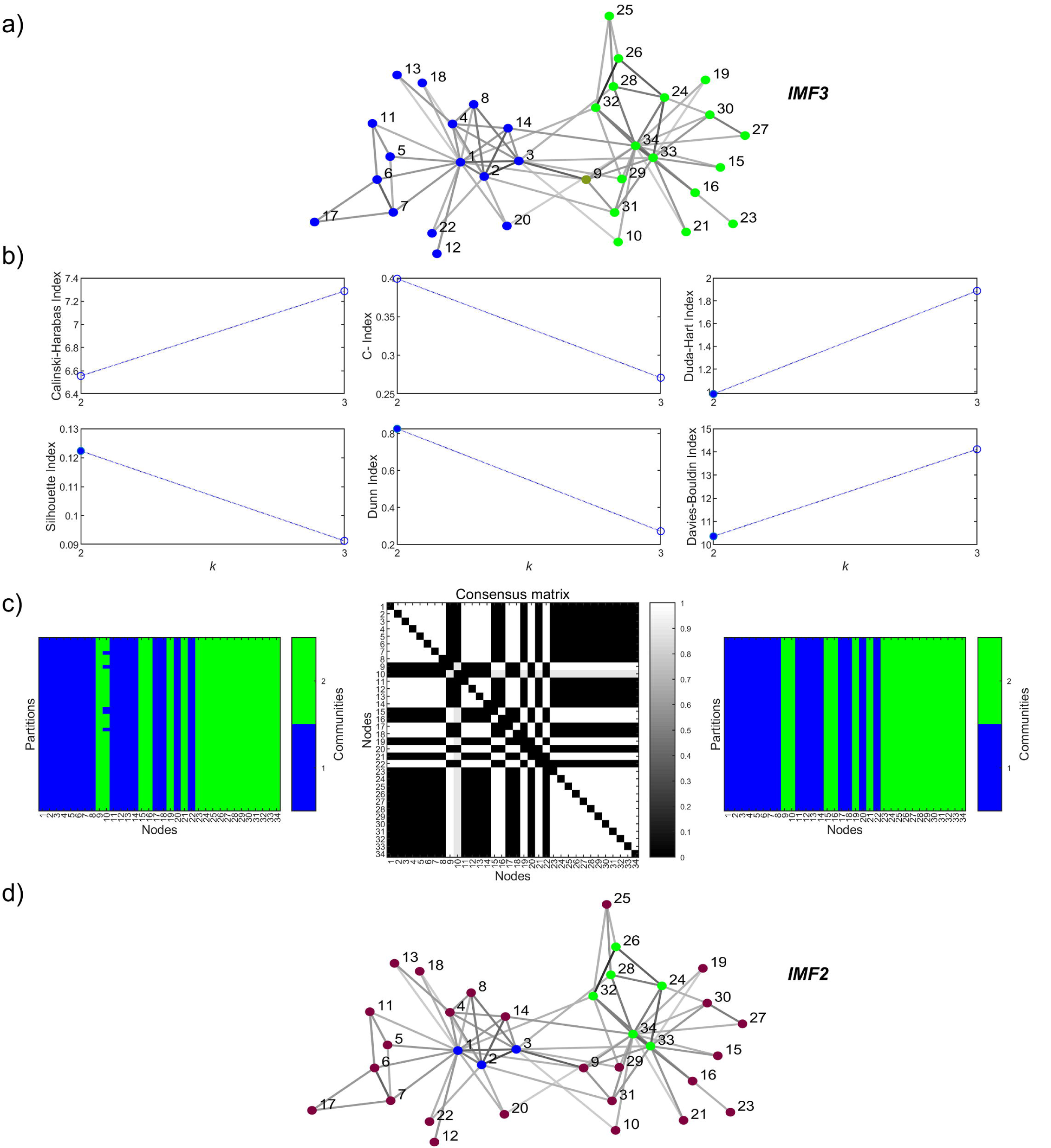
Communities of Zachary’s karate club. a) Representation of the community structure obtained over IMF_3_. The two groups in which the network split after the conflict largely coincide with this pattern. The instructor’s (president’s) faction is shown in blue (green). The node colored in olive is misclassified as belonging to the president’s faction when compared to the ground-truth. The edges drawn are proportional to the weights of the connections. b) Validation indexes supporting the selection of *k* = 2 clusters in the data corresponding to IMF_3_. Values of *k* for which *k*-modes yields singleton communities are not shown (*k* = [4, 20], *k* ∈ ℕ). c) Consensus clustering for partitions obtained with 50 different sets of random features of IMF_3_. d) Over IMF_2_, a new partition of three communities is obtained with small clusters including the instructor and the president.

The other structure (Fig. 4d) was obtained at IMF_2_. This consists of three communities and suggests a pattern in which the two leaders (node 1, the instructor, and 34, the president) often interact with what presumably is their intimate friendship circles (nodes colored in blue and green, respectively) and the rest of the network conforms a different group. It is important to bear in mind that our analysis was performed over a weighted matrix accounting for several contexts in which the members of the club were seen interacting. Thus, the broader community found may represent a set of passive actors in the fission of the social network, some who “sit and wait” for the inputs coming from the rapidly exchanging groups of leaders and close followers. Therefore, the consideration of temporal scales –essential to our methodology– could be a key aspect to uncover new and interesting phenomena.

Visualization of the community structures was achieved by means of SpringVisCom (Jeub, Balachandran, Porter, Mucha, & Mahoney, 2015).

### 3.2 Community detection in brain networks

#### 3.2.1 The macaque visual and sensorimotor network

After testing the reliability of our method in several networks for which the community structure is known, we proceeded to its application to brain graphs. The first network considered was that of binary connections between cortical structures of the macaque visual and somatosensory-motor systems (see *Methods, Macaque visual and sensorimotor anatomical network)*. As such, a certain distribution of network nodes between those two functional systems was expected. Fig. 5 shows the hierarchical tree returned as consensus clustering for the macaque anatomical network. At the highest level (two-clusters partition), the communities found correspond with the documented distinction between visual and sensorimotor areas (Hilgetag, Burns, O’Neill, Scannell, & Young, 2000; Van Essen & Felleman, 1991; M P Young, 1993). The sensorimotor system retained a single hierarchy, comprised of areas 3a, 3b, 1, 2, 5, Ri, S2, 7b, IG, ID, 4, 6 and SMA, at the following level whereas the other community split in two (showed in variations of blue). The first of the groups is composed of areas V1, V2, V3, VP, V3a, V4, V4t, MT, MSTd, MSTl, FST, PO, PIP, LIP, VIP and DP. The following cortical regions appear in the other set discovered: VOT, PITd, PITv, CITd, CITv, AITd, AITv, STPp, STPa, TF, TH, 7a, FEF, 46, TGV, ER and 35. These two smaller clusters largely resemble the traditional anatomical subdivision of the primate visual system in groups of ‘ventral’ and ‘dorsal’ areas (Hilgetag et al., 2000).

**Fig. 5.**
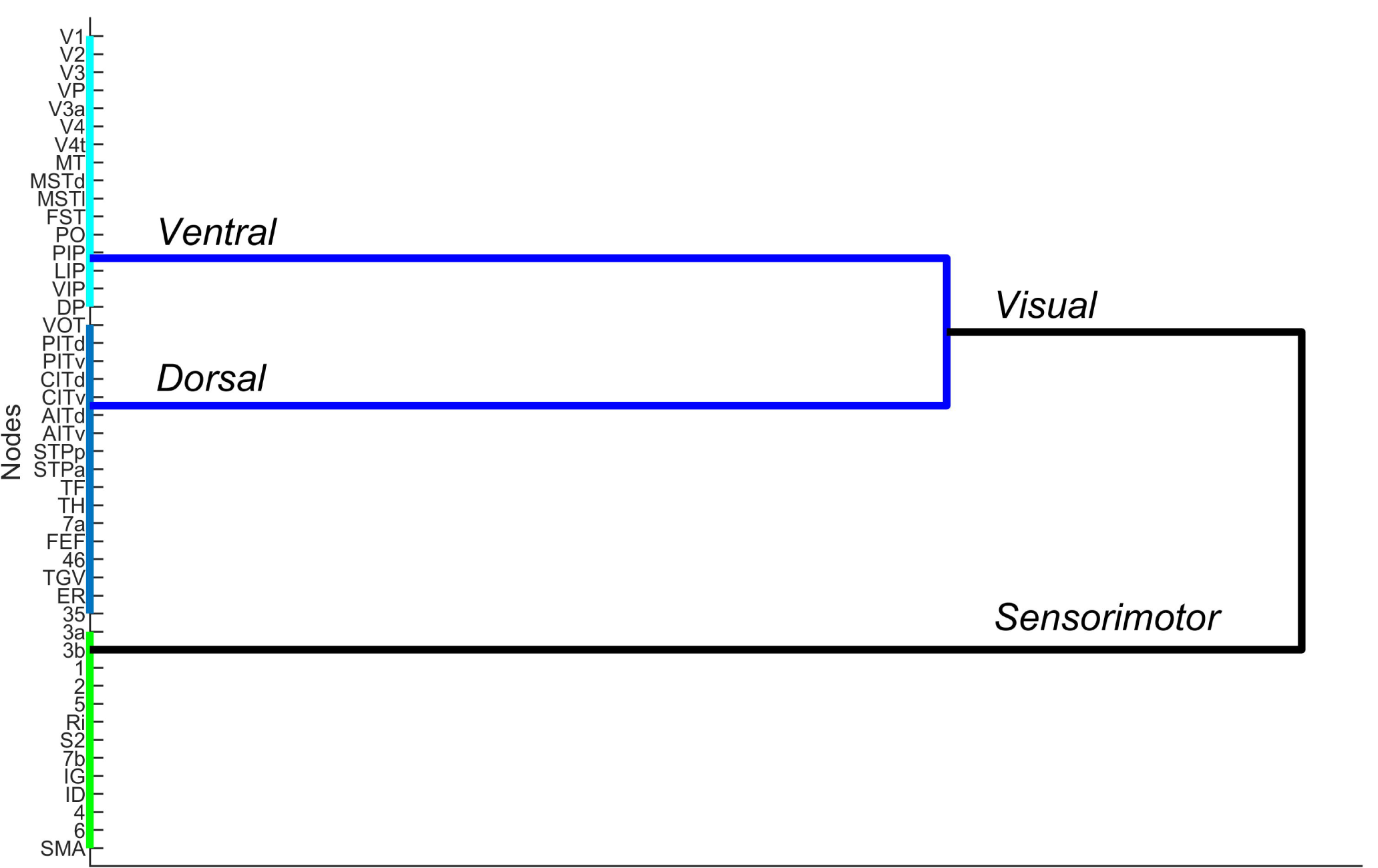
Dendrogram for the hierarchical consensus clustering of the primate visual and sensorimotor cortex. Somatosensory and motor areas are colored in green. Regions largely regarded as part of the visual system appear in blue. These are divided in two groups for predominantly dorsal and ventral anatomical areas.

To further explore the performance of the algorithm herein introduced, we compared our results to the more conventional Louvain-like community detection methods –Newman-Girvan null model, implemented in the Brain Connectivity Toolbox (Rubinov & Sporns, 2010). Fig. S5 shows the results of 50 initial runs and the consensus partitions obtained in both cases. Both widely used criteria for the selection of the resolution parameter of the Louvain algorithm, i.e., *γ* = 1 or *γ* chosen as the value for which partitions are more similar, yielded the same result, which is a three-clusters structure. This organizational pattern echoed our last result (Fig. 5, Fig. S5b), except for the ventral occipitotemporal (VOT) cortex which in some partitions appeared together with the ventral cortex though was eventually grouped with most dorsal areas.

#### 3.2.2 The human brain networks

We have also applied our community detection algorithm to the network of 78 cortical and subcortical neural conglomerates of the human brain (see *Methods, Human brain anatomical network* and *Simulated human brain functional network*). Fig. 6 summarizes the results. Firstly, the anatomical connectivity matrix was considered. We obtained two organizational levels, which are depicted in Fig. 6a and 6b. The highest of the two (Fig. 6a) consists of two communities which are the left and right hemispheres of the brain. Running the clustering algorithm with the features of a different IMF yielded a structure of subdivisions of the two hemispheres (Fig. 6b). This four-community organization is practically symmetrical except for the postcentral gyrus, the pallidum and the thalamus proper, which switch communities from one hemisphere to the other (see Fig. 6b and Table S2). The two communities to the top of the brain representation in the panel are mainly part of the frontal lobe, the cingulate cortex, and the basal ganglia. On the other hand, those shown toward the bottom generally correspond with parietal, occipital and temporal areas (Klein & Tourville, 2012; Lanciego, Luquin, & Obeso, 2012). An instance of the validation indexes supporting the existence of four communities in this data was given as a demonstrative example in Fig. 2c.

**Fig. 6.**
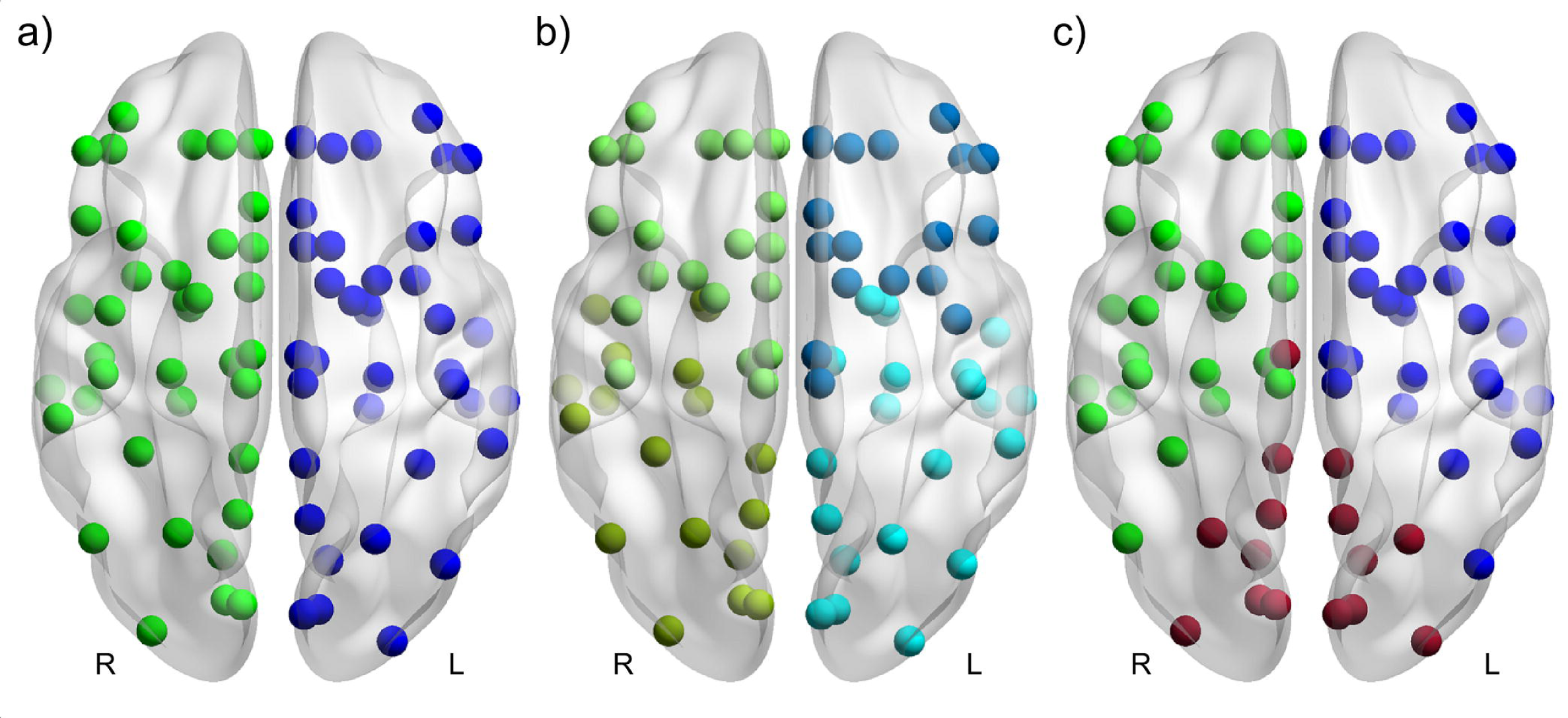
Representation of the communities of the 78-regions human brain anatomical and simulated functional networks. a) Two-communities structure obtained for the anatomical network. b) Four-communities structure obtained for the anatomical network. c) Three-communities structure obtained for the functional network with global coupling parameter of the Kuramoto oscillators *κ* = 5. Colored nodes correspond to communities and their location, to average coordinates of the brain regions in MNI space.

The three synthetic functional networks (Fig 3b-d) resulting from superimposing Kuramoto oscillators to the matrix of anatomical connections were explored afterward. In Fig. 6c, the only communities found in the *κ* = 5 case are shown. Two of those communities are a set of neural structures belonging to either the left (in blue) or right (in green) hemisphere. However, a third community (in maroon) consists of fifteen inter-hemispherical regions, all of which except for the right posterior cingulate appeared in a symmetrical manner in both hemispheres, including the totality of the occipital areas (see also Table S2). Fig. S6 shows the consensus clusters identified by using the two standard criteria for the resolution parameter in Louvain-modularity maximization. While three regions are grouped inter-hemispherically with those of their kind, in general, modularity maximization seems to fail at recognizing the functional relationships that are supposed to exist in this data, e.g., the mixed community pinpointed by our method. The tendency to split communities to obtain higher modularity values is also observed in Fig. S6 as anatomical communities are divided in a virtually arbitrary way (compare to Fig. 6b, for example). For *κ* = 30, our algorithm (and Louvain-maximization) returned the same two-hemispheres structure illustrated in Fig. 6a. Nevertheless, for *κ* = 150, no community structure was found over any of the IMFs and combinations of walkers considered (one cluster encompassed all nodes). This conclusion was reached by applying the criteria of *Appendix C*.

Visualization of the community structures was achieved by means of BrainNet Viewer (Xia, Wang, & He, 2013).

Lastly, community finding was applied to the connectivity matrix of the functionally specialized cortical regions resulting from the ICA analysis of the HCP500 “PTN” data (see *Methods, Human brain resting-state fMRI network)*. The main results are shown in Fig. 7. Five clusters were found (IMF_3_). We have labeled these communities according to their equivalence to conventional functional sub-systems observed in resting-state fMRI (see Table S1). For example, a mostly-visual cluster I included several visual regions, one dorsal attention (ICA map 8) and one somatomotor (ICA map 22) (Sacchet et al., 2016; Thomas Yeo et al., 2011). The ventral attention and a mix of the default mode network (DMN) and the somatomotor sub-system were also quasi-pinpointed at this temporal scale, whereas purely visual (II) and frontoparietal clusters appeared as well. Although not shown here, a different partition arose at a slowest IMF containing two clusters, one of them including the ICA maps 6, 11, 14 and 19, i.e. the cluster Visual II and members of DMN/Somatomotor.

**Fig. 7.**
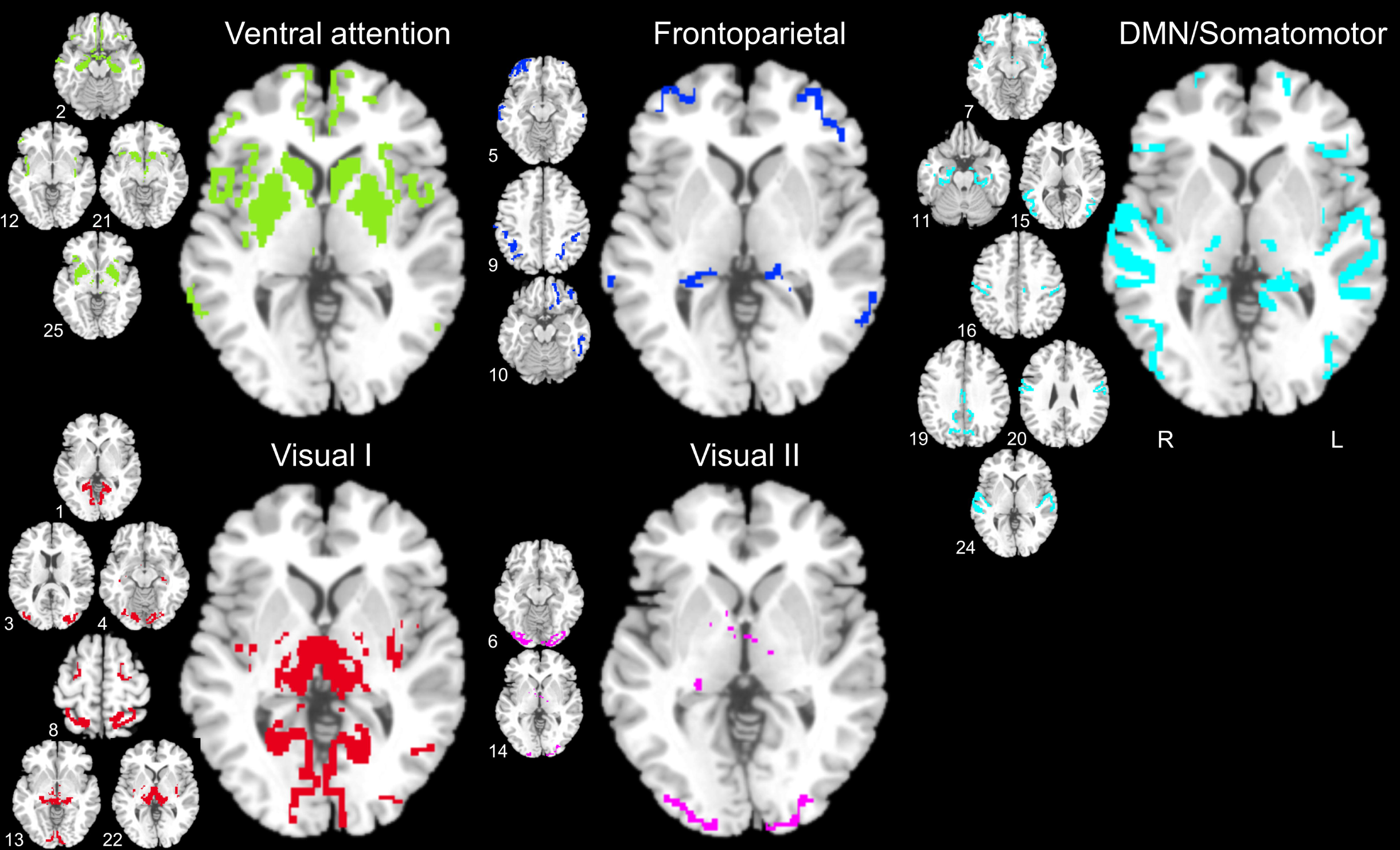
Clusters of the ICA-parcellated healthy human brain connectome. Five communities exist with a high degree of correspondence with well-established resting-state sub-systems (Ventral attention, in olive; Frontoparietal, in blue; DMN/Somatomotor, in cyan; Visual I, in red and Visual II, in violet). Representative axial slices of the spatial components (nodes) combining to form the clusters are shown next to them. For consistency, all the clusters are illustrated through views of their center axial slice.

The overlay views were created with MRIcron (https://www.nitrc.org/projects/mricron).

### 3.3 Interpreting the communities identified by the method

Given the novelty of the method, we consider opportune to provide an analysis –*a posteriori*– of the unveiled community structures and reiterate the checkpoints and good practices we recommend the end-user to implement, to retrieve meaningful communities. Let us start by illustrating the interrogation of the communities obtained over all the IMFs to retain only the meaningful patterns. This is explained in *Methods, Accepting/rejecting hierarchical partitions* and *Appendices B* and *C*. The set of codes accompanying the paper (see *Data and code availability statement*) also incorporates a user-friendly graphical representation of the acceptance/rejection guidelines. We show the process in Fig. S7 (exemplary first six IMFs of Zachary’s network for visualization purposes). As mentioned (*Results, Community detection in benchmark graphs*), IMF_3_ (2 communities) and IMF_2_ (3 communities) were chosen as a majority of the validation indices (*Appendix B*) pinpointed such patterns. Conversely, IMF_5_ and IMF_1_ were discarded as the *Davies-Bouldin* index’s decision rule (*Appendix C*) suggested that a full set of singleton communities was more likely to exist in the data than any organizational pattern. On the other hand, the patterns produced with the features of IMF_6_ and IMF_4_ were discarded because no majority was achieved.

Additionally, a two-communities pattern would have been replaced by the one obtained over IMF_3_ (Zachary’s ground-truth clusters). The main reason for this design choice is the expectation that slower time scales correspond to larger communities –and vice versa (Delvenne et al., 2010; Fortunato & Hric, 2016) –see below. Zachary’s network’s IMF_4_ also constitutes a rare example of noisy IMF in the sense of being a signal containing no more information than what pure noise does (Wu & Huang, 2004) (Fig. S8). In terms of statistical significance, the user may well reject the partitions obtained when the IMFs used to build the clustering features are not informative.

The application of these guidelines may ensure the correct selection of organizational patterns in the network. Nevertheless, a necessary condition for successfully selecting the communities existing in the generic *n*-dimensional network is guaranteeing that the fraction of walkers-signal does sample the network and the nodes associations. How many walkers are required for this? In *Methods, The network’s signal* we express *W* ≫ *n*. One good initial consideration would be having a number of walkers, *W*, at least 10 times bigger than the number of network nodes. We performed an analysis of the stability of the partitions returned while changing the relationship 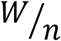 in two networks: Zachary’s karate club of 34 nodes and the human brain-anatomical, with 78 (see Fig. S9). For these two real networks –and not for the rest– we had prior knowledge of the ground-truth partitions, i.e. the two fractions in which the karate club separated (obtained at its IMF_3_) and the expectation that the two hemispheres of the brain should be separated by the community detection method (which occurred at its IMF_6_). Therefore, adjusted mutual information can be utilized to provide a measure of the robustness of the method at returning these partitions while 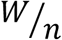 is increased from its minimum possible value (the 200 walkers that were selected to build the clustering features). In the range up to 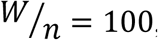, the communities returned by the method generally match the ground-truth communities (with the exception of the above-mentioned node 9 of Zachary’s network thus yielding an AMI of ∼0.83). In effect, the initial assumption of *W* > 10 · *n* which made us fix *W* = 1000 for all networks in this study, seems to suffice. For other networks, particularly those of neuroimaging applications, the user could extrapolate the information of Fig. S9 to minimize computing time and consider the characteristics of his/her networks, namely the number of nodes, the density of connections, etc. On this note, how effective are our clustering features in representing the topology of the networks? While grouping nodes by their co-appearances between zero-crossings of the IMFs, nodes with unusually high connectivity (hubs) appear more often than less connected nodes do. This is shown in Fig. S10 where the probability of finding a node between zero-crossings is represented as a function of the strength (“weighted degree”) of the connections that such a node has (Rubinov & Sporns, 2010; Sanchez-Rodriguez et al., 2018) for IMF_2_ of Zachary’s network. Over this IMF, three communities were identified. Two of them included the highly connected nodes 1 and 34 and their close ‘cliques’, while the remaining cluster was formed by somewhat less connected nodes (in maroon). However, this cluster is not exclusively made up of low-strength nodes, with at least six members being more connected than others in the leaders’ cliques. Topologically, connections exist for the movement of walkers between the nodes in the community of the ‘passive actors’ (distinct of the leaders and their closest followers), determining the dynamics of the social network. The highly symmetrical four communities of the human brain anatomical network (IMF_4_) also present a heterogeneous distribution of node strengths. The hubs of this network, corresponding to the mean + 1 SD of the strength distribution (Fulcher & Fornito, 2016), are not equally present in all communities. Four of them appear in the ‘Left parietal-occipital-temporal’ community (L lateral occipital, L precuneus, L superior temporal, L hippocampus), two in ‘Left frontal-cingulate-basal’ (L superior frontal, L insula), two in ‘Right parietal-occipital-temporal’ (R lateral occipital, R superior temporal) and only one in ‘Right frontal-cingulate-basal’ (R superior frontal). It is the network’s topology (thus: nodes appearing together between our zero-crossing features of the IMFs) and not the mere node strength-dependent probability of finding a node in the features what defines the organizational patterns that were identified. If anything, these examples hint at the very essence of our method: the network signal and features sample the information flow on the considered network. The information flow is related to the topological properties of the network (Sotero et al., 2019).

Then, what physical (and biological) meaning do the communities obtained by our method have? What is the exact relationship between the IMFs (temporal scales) of the network’s signal and the spatial scales of the obtained communities? To clarify these issues, we considered all the real networks herein studied, their hierarchical patterns, and the IMFs at which these meaningful (see above) communities appeared. Then, for each of the reported communities, we calculated the (within-community) characteristic path length (Rubinov & Sporns, 2010; Sanchez-Rodriguez et al., 2018), i.e. the mean shortest path length between all pairs of nodes belonging to that community. The overall pattern is spatially characterized by the average characteristic path length taken across all the communities in the pattern, divided by the full network’s characteristic path length, a measure we term ‘average normalized within-communities path length’. Take, for example, the human brain anatomical network and its four communities of IMF_4_. Each of these clusters has characteristic path lengths of 1.95, 2.15, 2.02, and 1.99, while the network of 78 nodes, 3.37. The normalized within-communities path length of approximately 0.60 thus defines the expected shortest distance that a walker travels to visit the nodes pertaining to one of the communities formed compared to the mean shortest distance it travels to visit any network node. On the other hand, the IMFs per se represent the frequencies at which the information flow inside those communities occurs. Since these can be somewhat irregular signals (see Fig. 1), we used the average of the instantaneous frequencies, calculated with MATLAB’s *instfreq.m*. Here, we assumed a sampling rate of 1*Hz* for all networks with the purpose of illustrating their behavior in one single chart, even though this sampling rate value lacks significance in terms of the propagation of the walkers within each network. Finally, a grand average over the walkers was performed as all of them contributed to the clustering features that yielded the reported community structures. The results are shown in Fig. 8. It is important to bear in mind that through the above-mentioned validation indexes and decision rules, no more than a couple of community structures were identified for the considered communities, thus rendering impossible to rigorously fit the curves in Fig. 8 to a mathematical law. However, seeing each network separately –as should be, given that they are independent entities– one can notice a 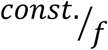-behavior of the average within-communities path length. In other words, faster time scales (higher frequencies of the IMFs) correspond to shorter effective distances in the communities. The constant in these relationships may define the velocity of the propagation of the walkers, or information flow, in each network.

**Fig. 8.**
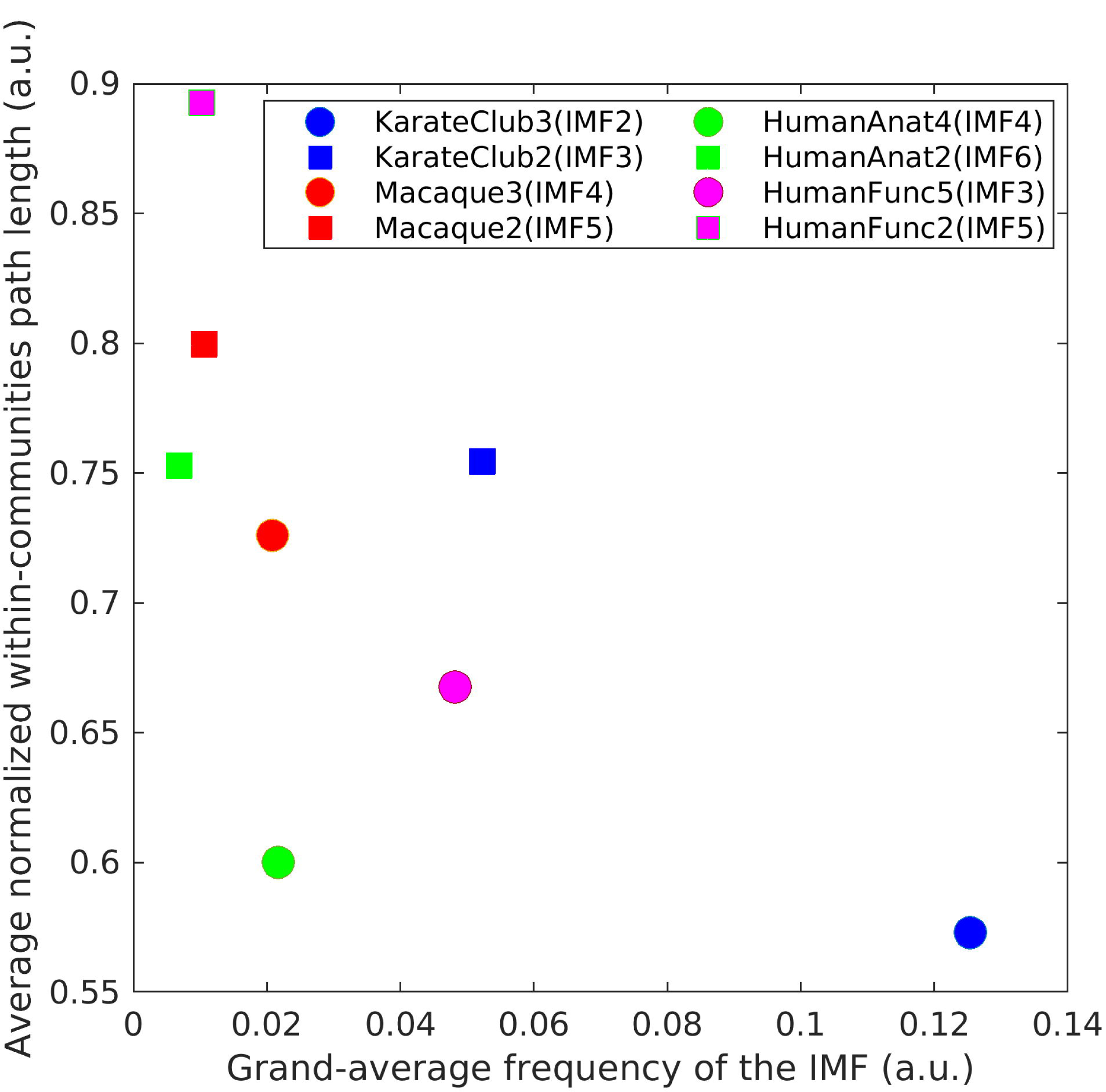
Relationships between spatial and temporal scales of the real networks considered in this work. Each network has been identified with a different color. The clustering features corresponding to two IMFs yielded community structure in each case. The number of communities existing at these IMFs are written next to the networks’ names (KarateClub: Zachary’s karate club network; Macaque: macaque visual and sensorimotor network; HumanAnat: human brain anatomical network, HumanFunc: Human brain resting-state fMRI network). Circles represent the fastest IMFs, while squares are used for the slowest ones, in each case. Temporal scales are characterized in terms of a grand-average frequency, by taking the mean value of the IMF’s instantaneous frequencies over all the walkers and assuming a sampling rate of 1Hz for visualization purposes. The spatial scales are described by the average shortest path length of the nodes corresponding to one community, divided by the overall network’s characteristic path length.

## 4. Discussion

The problem of identifying modular structures at different scales of a network has captured the attention of the neuroscience community in recent times. Notably, Jeub et al. (Jeub et al., 2018) and Ashourvan et al. (Ashourvan et al., 2019) have introduced variants for the sweep through the Newman-Girvan modularity’s γ-space eventually yielding hierarchical architecture. These methods have been tested in brain networks with encouraging results. Inherent limitations exist, however, as the algorithms build on multi-scale modularity functions. Consequently, the exposed structures depend on the selection of several parameters and a null model, as in regular Louvain-like community detection (Blondel et al., 2008). Specifically, authors tend to recommend the utilization of null models that suit the characteristics of the data perfectly (Betzel et al., 2017). However, null models appear as abundant as detection algorithms in the literature oftentimes, making its selection a key step for the success or failure of the application of an algorithm (Sporns, 2013). Ideally, one would like to provide the user with minimum-input tools that can reveal the underlying structures of the data in natural ways.

In the recent past, much of the discussion as to the directions of neuroscience research was centered on avoiding univariate statistical comparisons and, instead, looking at the network interactions as a whole (Telesford, Simpson, Burdette, Hayasaka, & Laurienti, 2011). What is more, we believe that to better capture the complexities of a system like the brain, with multiple spatio-temporal scales and dynamic reconfigurations, the mere application of generic network science methods is not sufficient. Tools must be developed to account for the brain’s unique characteristics. Thus, in this paper, we have searched for organizational hierarchies through the temporal scales of the network’s random walker signals, without necessitating to fix any model parameter. By doing so, the characteristics of the information flow in the brain are also incorporated (Sotero et al., 2019). The integration of the network’s architecture with the dynamical interactions of the oscillatory modes in the large-scale brain is consequently suggested as an important factor in clustering techniques. Likewise, long running times (Nguyen, 2017) and a finite number of IMFs in the signal decomposition to investigate the existence of communities (N. E. Huang et al., 1996), could render the algorithm ill-suited for applications on high-dimensional graphs (different than large-scale neuroimaging-based networks). In that sense, other random walkers-based clustering techniques, including Walktrap (Pons & Latapy, 2005), Infomap (M. Rosvall & Bergstrom, 2008), and the method by (Delvenne et al., 2010), have successfully dealt with the demands of clustering large networks. In particular, Delvenne et al. (Delvenne et al., 2010) explored network time scales through the autocovariance of the clustered Markov process associated with the random walk. Natural communities persist over time, which is reflected in the autocovariance. This elegant approach was proved efficient in binary undirected graphs of various topological characteristics and sizes. To the best of our knowledge, applications in the scope of neuroimaging were not demonstrated.

### 4.1 The brain organizes according to function

In discussing the results of applying our methodology to the macaque visual and sensorimotor network, several insights can be gleaned. Firstly, the obvious and most straightforward precedent is the classification of areas according to the functional neural system they belong to, visual and sensorimotor (Van Essen & Felleman, 1991; M P Young, 1993). Additionally, two anatomical pathways have been identified in the visual system (Mishkin, Ungerleider, & Macko, 1983), which are usually known as dorsal (originating in the occipital cortex and terminating in the parietal lobe) and ventral (from occipital to temporal). These anatomically constrained divisions constituted the rationale for expecting the separation between somatosensory-motor and visual areas (which in turn was further divided into dorsal and ventral) by our community detection algorithm, given the numerous connections existing between areas in a functional system and somehow less connections with outsiders (Christopher J. Honey, Thivierge, & Sporns, 2010). We highlight, however, that limbic structures like the entorhinal cortex (ER) and the perirhinal cortex (A35), usually considered together with the sensorimotor system (Van Essen & Felleman, 1991), were clustered with most visual areas. Also, the VOT appeared in an otherwise dorsal community. This result is analogous to the one described by Hilgetag et al. (Hilgetag et al., 2000) in a study on cluster organization of a similar, larger network using now obsolete techniques. Moreover, in that work, prototypical ventral and dorsal areas V4 and 7a were clustered with the opposite streams. Notwithstanding the slight differences, the subdivision of the visual community obtained here closely resembles the ventral and dorsal streams reported by Hilgetag et al. Several more recent studies utilizing modularity maximization methods have detected similar sets of communities, yet different macaque datasets have been used (Sporns & Betzel, 2016).

We believe that the minor discrepancies in the analysis of the primate cortical network commented in the above paragraph are due to two main issues. First, we should revisit the limitations that exist intrinsic to each community detection method. These, of course, affect the partitions returned given any connectivity matrix, revealing the necessity of continuing to develop tools and migrating to more comprehensive approaches that use as much valuable network information as possible. Another important matter to recall is that imperfections exist in data as well. For example, the matrix of connections we used (M P Young, 1993) encompasses reports from several studies, which sometimes even employed different anatomical parcellations. Also, this matrix accounts only for the existence or absence of reports of links between areas of the macaque cortex, without considering the strength of the connections. Whether datasets accurately reflect the particularities of the connections existing in the brain or not will remain a fundamental question in neuroscience.

The next application of our newly introduced community-finding method was to human brain networks. We considered a connectivity matrix in which each entry reflects the evidence of the existence of a white matter link between two brain regions (Y. Iturria-Medina et al., 2007), given a template of such connections in the young healthy brain (Dunovan et al., 2015). Dominant connections were retained through a minimum spanning tree algorithm. Although the minimum spanning tree trims connections and does not contain loops, it is believed to provide a correct representation of any denser brain network to which it is applied, retaining paramount topological characteristics like its small-worldness and scale-freeness (Tewarie, van Dellen, Hillebrand, & Stam, 2015).

We have found two partitions of the anatomical network, which seemingly follow physical proximity and functional specialization rules. In the case of the first partition, commissural fibers appear to act as those rare links to members of other communities for the two brain hemispheres were perfectly separated. Each hemisphere split into two communities over the other partition found. The four-communities structure was almost bilaterally symmetrical, as only the postcentral gyrus, pallidum and thalamus proper exchanged membership, i.e., they grouped with most frontal, cingulate and basal ganglia regions in the right hemisphere and with parietal-occipital-temporal areas in the left. Variations in local connections within hemispheres may be the reason why these regions behaved in such a way. In principle, the thalamus, as universal relay station, and the pallidum projecting to the thalamus (Lanciego et al., 2012) should have no constraints to belong to one or the other intra-hemisphere community found. The postcentral gyrus, although deemed part of the parietal lobe is in the vicinity of the frontal lobe, possibly explaining its grouping with such neural structures. The association of brain regions to perform processes and functions could also be reflected in the anatomical network and, consequently, in the communities obtained. For instance, having the basal forebrain clustered with frontal areas may be justified by the fact that its projections to the prefrontal cortex are paramount for attention, learning and memory, and decision-making (Tashakori-Sabzevar & Ward, 2018).

The study of the matrices for the interaction of oscillators over the anatomical frame yielded stimulating results. For one thing, the mechanism for the transitions between functional states relates to the tuning of the coupling parameter in the model. Three communities appeared in the low-coupling regime (*κ* = 5), one of them presenting areas from both brain hemispheres in a close-to-symmetrical pattern. When the coupling strength was raised to *κ* = 30, the mixed community was destroyed and the only recognizable pattern was the one of two separate brain hemispheres, which was one already existing in the anatomical network. This is because the connectivity matrix of a set of Kuramoto oscillators overcomes the dispersion of natural frequencies for higher values of the coupling parameter (Breakspear et al., 2010). Higher functional couplings amplify the anatomical subdivision of the network in two hemispheres.

Other characteristics of the clusters of the simulated functional networks are also noteworthy. For example, all the occipital areas (Klein & Tourville, 2012), responsible for vision (Johns, 2014), appeared in the mixed community of the *κ* = 5 -regime. Previous studies of modular organization in functional graphs found robust grouping in the occipital cortex (Meunier, 2009). Among the other areas present in the intra-hemispherical cluster: the superior parietal lobe has abundant connections with the occipital lobe and participates in visuospatial perception (Johns, 2014); the precuneus has a major role in visuo-spatial imagery (Cavanna & Trimble, 2006) and the posterior cingulate cortex is considered a core node of the DMN and to be involved in many tasks (Yasser Iturria-Medina et al., 2014). The inter-hemispherical community obtained appears to be one extended circuit concerned with the function of vision. The rest of the nodes within each hemisphere may consequently process all the non-visual stimuli, possibly constituting an optimal configuration for speedy and accurate performance on cognitive tasks (Garcia et al., 2018). We believe that the detection of these communities supports the notion of functional integration in the brain, whereas evidence for segregation can also be found in a division that isolates units specialized in handling with visual stimuli.

Although a valuable exploratory tool, *in-silico* brain functional networks do not totally reflect the vast amalgam of phenomena present in real data. Motivated by this concern, we proceeded to study empirical resting-state fMRI data in the healthy brain. Out of the many possible options for defining fMRI connectivity that are currently used by the neuroscience community (Bassett et al., 2013; Bordier et al., 2017; Moradi et al., 2019; Smith et al., 2013), we decided to inherit the connectivity estimations performed by the HCP on the dataset we used (Smith et al., 2013, 2015). The approach of the HCP builds on the utilization of the data-driven ICA algorithm (Mckeown et al., 1998) to simultaneously obtain multiple spatial maps, each of them having relatively homogeneous connectivity profiles and functional specialization (Smith et al., 2013). In agreement with the rest of the situations studied in this paper, which are in turn inspired by classic network neuroscience methodologies (Bassett et al., 2013; Betzel et al., 2017; Christopher J Honey et al., 2007), we treated the functional regions as non-overlapping though these regions are poly-functional in essence (Betzel et al., 2016). The straight-forward upgrade that should be considered in future analyses to allow for the existence of overlapping communities –thus recognizing functional multiplicity– is replacing the “hard” clustering *k*-modes technique with its fuzzy counterpart (Z. Huang & Ng, 1999). Additionally, despite the existence of finer ICA-parcellations, we adopted a coarse subdivision to seek a (nearly) one-to-one relationship of the nodes in the connectivity matrix with studies that replicate systems of functionally coupled regions across the cerebral cortex (Thomas Yeo et al., 2011).

The main partition obtained significantly resembled the conventional subdivision of resting-state fMRI in functional systems. The visual cortex, once more, presented distinctive community patterns. It was further divided into two clusters. The smallest of the two (Visual II) only contained two dorsal ICA activation maps of the visual system. Moreover, one dorsal attention ICA map (8) was clustered with the mostly-visual community I, in line with evidence suggesting an interaction between these two functional sub-systems (Thomas Yeo et al., 2011). Another dorsal attention component was associated with the partly somatomotor cluster. Researchers have referred to the dorsal attention system as the prototype distributed cortical network (Thomas Yeo et al., 2011) given its strong functional coupling with sensory and motor regions. In previous work (Smith et al., 2013, 2015), the HCP applied hard agglomerative hierarchical clustering techniques implemented in MATLAB to a similar connectivity matrix. Despite the difficulties of their methodology for identifying meaningful partitions and the correct number of clusters, a large-scale pattern was observed, which mostly differentiated visual, sensory-motor and dorsal attention regions from cognitive-processing ones. Compared to these studies, our community detection algorithm yielded suboptimal results when producing a community of mostly DMN and somatomotor ICA maps together. This apparent discrepancy may be more closely related to the interpretation paradigm we adopted instead of the community detection *per se*. For example, the ICA map 24 was uniquely labeled as “Somatomotor 2” (Table S1), “Somatomotor 2” a system with which ICA map 24’s voxels had 59.5% overlap, ignoring altogether the 28.0% overlap it had with the system “DMN 4”. This is in strong contrast to, e.g., ICA map 6 which was 86.4% “Visual 1”. Thus, it is believed that ICA map 24’s connections somewhat had served as strong link between DMN and somatomotor regions of the studied parcellation, provoking their combined clustering. As above-stated, limitations of this sort could be overcome by considering the more realistic overlapping sense of resting-state functional connectivity.

Regarding the relationship between spatial scales and the IMFs, one may wonder how to interpret these temporal scales of the information flow in neuroimaging applications. For example, the temporal scales identified in functional neuroimaging data (Yuen, Osachoff, & Chen, 2019) are not to be confused with the signal decomposition we adopt in this study, because this, unlike the region of interest-oriented conventional neuroimaging analysis, is built from the random walker’s frame of reference, i.e. the fraction of other walkers that each one finds throughout its journey within the networks (see *Methods, The network signal*). Although it is possible to construct signals for the nodes (e.g., the number of walkers visiting at each time iteration), they would be altogether different than the ones we conceived. Consequently, the assumptions adopted and physical interpretation in terms of information flow would no longer stand. In the future, it will be interesting to design other forms of network signals to investigate relationships between measures derived from structural connectivity and functional connectivity as it has been done in the outstanding works of (Abdelnour, Voss, & Raj, 2014; Chiêm, Crevecoeur, & Delvenne, 2020; C J Honey et al., 2009). For the moment, the analysis of functional data and its topologically peculiar networks (Gates et al., 2016) through our algorithm must be plainly seen as the application of a reliable community detection method (see *Results, The human brain networks* where real resting-state sub-systems and simulated functional clusters were identified), thus avoiding the temptation to overinterpret multi-scale functional relationships. As a proof of concept only, we can calculate the IMFs grand-average frequencies (Fig. 8) assuming a sampling time of 1/*TR*, as the *TR* defines the resolution of the fMRI data, thus indirectly affecting the connectivity matrix and the propagation of walkers in it (*TR* = 0.78*s* (Smith et al., 2015)). In this case, the frequencies of the IMFs whose features return validated communities are inside the interval corresponding to the low-frequency (0.01–0.08 Hz) fluctuations in fMRI (Biswal, Kylen, & Hyde, 1997; Zou et al., 2008) while faster and slower IMFs than these two had frequencies outside that range. Structural connections, on the other hand, constitute the physical scaffold over which interactions occur in the brain, which perfectly matches the notion of information flow. For anatomical neuroimaging networks, we can then straightforwardly relate the IMFs of the network signal with neuronal electrical signaling just by assuming that the motion of the random walkers represents electrical impulses traveling through the brain. The sampling time in the analysis yielding to Fig. 8 can be replaced by axonal conduction delays (other delays being neglected for the sake of simplicity): we record the walker’s IMFs each time it has landed in the following brain region of its path. While conduction delays depend on several factors that influence axonal conduction velocity and on the distance between the involved sites, we fixed a 7*ms* universal value, this being one of the typical human callosal conduction delays reported by (Caminiti et al., 2013). Then, the IMFs (6 and 4) detecting communities in the human brain anatomical network had grand-average frequencies of approximately 0.98 and 3.10 Hz. In other words, the global oscillations separating the anatomical network in two and four communities present frequencies that coincidently lie in the range of the EEG delta waves. It has been hypothesized that sustained delta oscillations inhibit the activity of neural sub-systems that interfere with accomplishing certain tasks (Harmony, 2013). If this were the case, one could explain transitions from one processing scale to another (with their respective associated brain organizational patterns) as the network’s re-arrangement to successfully accomplish tasks. Further investigation could help clarify physiological mechanisms of network multi-scaling in health and disease (Sanchez-Rodriguez et al., 2018) by using realistic measures derived from our information flow-centered method.

### 4.2 On the strengths and limitations of the algorithm

To conclude this section, we would like to highlight important features of our method. One interesting scenario is the one of functional interactions with *κ* = 150. The large synchronization seen there yields a close-to-*n*-regular graph (Fig. 3d) that goes together with the algorithm identifying a single group of nodes. In fact, one recommended practice for testing new community detection algorithms is checking that they do not return group structure in the absence of it (Hric et al., 2014). Of further value is the effective performance shown when searching for the community structure of the benchmark graphs, all of which presented different network characteristics. On this topic, one must also mention the diversity of the real networks whose organization in groups was explored. The karate club is a small, weighted and undirected network. On the other hand, the macaque visual and sensorimotor network is binary and directed. The human brain networks, most of them with larger dimensions, had different levels of sparsity. Many algorithms, e.g. Infomap and (Delvenne et al., 2010) are initially designed for a specific type of graph (Fortunato & Hric, 2016) and only later extended. However, our method seems to be primed to perform reliable community detection in moderate-size networks like the ones associated with the brain’s large-scale activity.

It was not in our interest, though, to test its suitability for high-dimensional graphs. In that case, the demand for computing resources would grow. Firstly, the computational cost of *k*-modes scales linearly with the number of objects and many random initializations of the modes are required to find a reliable clustering solution (Nguyen, 2017). Secondly, the number of possible combinations of nodes appearing between zero-crossings of an IMF would increase as well, so the implementation of *k*-modes must be optimal to handle a large number of binary features. We bypassed some of these complications by using the University of Calgary’s (https://hpc.ucalgary.ca/resources) and Compute Canada’s (https://docs.computecanada.ca/wiki/Getting_started) computing clusters resources, where calculations were run in parallel. The other two stages of the algorithm, namely the random walks and empirical mode decomposition are already fast enough through built-in functions in MATLAB. Other speeding-up alternatives for the clustering problem must be explored, however. In the case of functional networks, we have shown how dimensionality reduction could be achieved by preceding the application of our EMD-*k-*modes algorithm with ICA-parcellation of the brain voxels, instead of trying to cluster the set of single voxels or highly granular parcellations. By doing so, we also suggest alternatives to integrate our method with existing functional clustering approaches and study hierarchical organization in high-dimensional problems.

Another solution for reducing dimensionality is the selection of relevant features in the data (Ronan et al., 2016). Because the random choice of features (i.e., the set of walks) could also generate solutions that differ from the existing structure if many unrepresentative features were combined, the identification of a relevant set is also desired for the stability of the method. Dimensionality reduction can be mainly achieved by feature reduction or feature selection. In the first approach, the variables are projected into a new space with a lower dimension (Alelyani Salem, Jiliang Tang, & Huan Liu., 2013). PCA and, more importantly, its categorical features-analogy, multiple correspondence analysis (MCA) (Greenacre & Blasius), retain the new features that capture the largest amount of covariance in the data (Ronan et al., 2016). However, these features cannot be linked to the original space and the application of PCA (MCA) to clustering problems oftentimes degrade the results if clusters appear in different subspaces (Lu et al., 2005). On the other hand, the selection of features that minimize redundancy is superior to feature reduction in terms of interpretability (Alelyani Salem et al., 2013) and performance (Ronan et al., 2016). Problems like the one our method is concerned with, binary feature selection for clustering, have rarely been addressed though, while most of the studies have focussed on numerical variables (Silvestre, Cardoso, & Figueiredo, 2015). To our knowledge, only a handful of works did explore clustering in the presence of categorical (thus also binary, in particular) data (Bontemps & Toussile, 2013; Silvestre et al., 2015). The methods therein developed make certain assumptions on the data and only solve the feature selection problem by simultaneously targeting a distribution in the desired number of clusters, which would not straightforwardly align with the rest of the pipeline in our algorithm. A desirable feature selection step would reduce dimensionality immediately before the application of *k-*modes. Dimensionality reduction by feature selection is a complicated matter on its own (Steinbach et al., 2004) and innovative solutions should be investigated. In short, we do not recommend the use of the method presented in this paper on large networks until further steps towards optimization are taken.

As a final note, we would like to discuss some concerns and recommendations that emerged from the initial utilizations of the algorithm. One issue is that no more than two hierarchical levels were identified over the IMFs of the networks analyzed in this study. Conversely, traditional methods like modularity maximization continuously divide clusters to satisfy mathematical constrains, which has also been pinpointed as a major limitation (see above and e.g. (Fortunato & Hric, 2016)). In lieu of accepting the network partitions returned by any method as the absolute truth without validation, the utilization of a broad set of metrics like the ones we define in *Appendix B* has been recommended (Bassett et al., 2013; Fortunato & Hric, 2016; Ronan et al., 2016). We can only know the real divisions existing in a handful of networks, like the simulated benchmark graphs and the karate club that divided into two fractions after a conflict (see *Results, Community detection in benchmark graphs*). Any other subdivision existing in, say the karate club, is only the product of the application of an algorithm and thus, ‘lives in that universe’. For the brain networks, the analysis is necessarily qualitative and based on previous anatomical-functional evidence (Klein & Tourville, 2012; Thomas Yeo et al., 2011), as it is obviously impossible to ascertain an absolute ground-truth (perhaps other than that the existence of two separate brain hemispheres in the vertebrate brain). If we define sparsity as 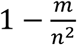 (Sotero et al., 2019) where *m* is the number of non-zero links and *n*, as usual, is the number of nodes, this value is higher for very sparse networks. With fewer connections, further divisions in communities are intuitively less likely. Table S3 shows sparsity values for several of the networks herein considered. The networks with known ground-truth partitions, LFR (100 nodes, 0.87 sparsity and 5 communities) and Zachary’s (34 nodes, 0.87 sparsity and 2 communities) thus offer a reference for the rest of the networks. Naturally, a dense network (0.16 sparsity) like the human brain functional connections of 22 nodes was identified with 5 communities, while the sparser (0.98) anatomical connections of 78 regions had 4 clusters at the finest level returning meaningful community structure in this network (IMF_4_) and no further subdivisions. This intuitive-critical analysis may serve to corroborate the suitability of the lowest hierarchical levels obtained in this work.

The second of the issues signaled by some users relates to the adoption of definitive consensus partitions to report when, for example (Fig. S5b) the VOT appeared clustered with most ventral areas in certain runs although being assigned to the dorsal community in the final partition. We understand that neglecting altogether some observed community associations (e.g. the VOT with the rest of the ventral areas) may be undesired in certain applications, as they could suggest a predefined operational flexibility of the brain areas to work in cooperation with others as in task-based functional connectivity (Gonzalez-Castillo & Bandettini, 2018). Similarly, our implementation ignores partitions that arise of the analysis of slower IMFs when the same number of communities is indicated at a faster IMF. In *Results, Interpreting the communities identified by the method*, we showed the inverse relationship existing between the average frequency of the IMFs and the spatial separateness of the nodes in the related communities (as given by the characteristic path lengths in the obtained clusters). Thus, the same number of communities identified with the features of a faster IMF are intuitively more likely to represent the preferred organization of the network in such a number of clusters. Although cases like this were not common in the networks analyzed, partitions rejected by the hierarchical condition could, again, be meaningful in scenarios of functional adaptability. Ultimately, the end-user will decide how to better employ the independent parts of our pipeline to satisfy their research question.

## 5. Conclusion

In summary, we have introduced an approach for the detection of modular organization by considering the temporal scales of the information flow over the networks of interest. This new tool may be particularly useful for the analysis of large-scale brain graphs, for which: 1) the transmission of information is a process of paramount importance and 2) a desirable balance between accuracy and computational complexity of the community detection algorithm can be achieved given the current implementation state. We find several organizational patterns existing in the brain anatomical and functional networks –also in the social network that we study. These structures may coexist together, in a dynamical way that is given by the temporal scales of the activity they produce, guaranteeing functional independence and coordination. Our results promise a shift of focus in the discussion surrounding the occurrence of community structure.

## Appendix A. Supplementary material

Supplementary data related to this article can be found at [*insert link here*].

## Appendix B. Cluster validation indexes

In what follows, *n* is the number of objects to be clustered and *k* is the number of such clusters. Let {*C*_1_, *C*_2_, *…, C*_*k*_} be a partition of the integers from 1 to *n* such that *i∊C*_*q*_ if the *i*th object,

***x***_*i*_, belongs to the *q*th cluster. The centroid (mode) of a subset *C*_*q*_ of *n*_*q*_ objects is the vector ***m***_*q*_ that minimizes the sum of the distances to all the objects in *C*_*q*_. Supposing the (mismatching similarity) distances between every *i* and *j, d*_*ij*_ are known, which also applies for the distances between objects and their clusters’ centroids, then:

-The *Calinski-Harabasz* index (Calinski & Harabasz, 1974), also known as pseudo-F ratio,

is defined as:

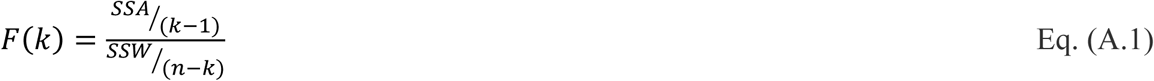

SSW and SSA are the within-group sum of squares and the among-group sum of squares, respectively. Adapting (Anderson, 2001), these quantities are obtained from the matrix of distances between pair of objects:

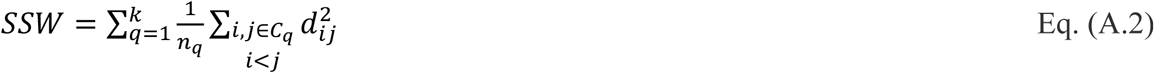

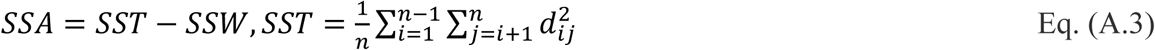

The among-group distances are large compared to the within-group distances in the case of high separateness and compactness. Thus, maximum values are taken to represent the correct number of clusters (Milligan & Cooper, 1985).

-The *C-Index* (Milligan & Cooper, 1985) is calculated as:

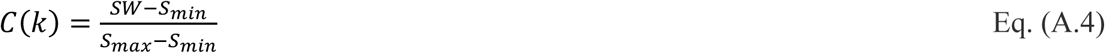

where *SW* is the sum of the within-cluster distances:

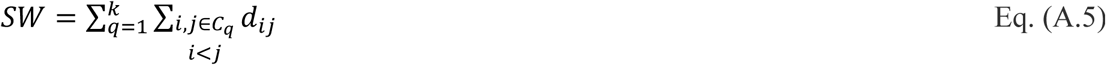

and *S*_*min*_(*S*_*max*_) is the sum of the *N*_*W*_ smallest (largest) distances in the dataset. *N*_*W*_ is the total number of pairs of objects in the same cluster, 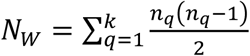 (Charrad et al., 2014). C-Index is restricted to the interval (0,1) and its minimum value suggests the optimal number of clusters (Milligan & Cooper, 1985).

-The *Duda-Hart* (Duda et al., 2001) score is inspired by the fact that the sum of squared-errors corresponding to a partition decreases with *k*. Thus, in conventional Euclidean-distance clustering problems, the optimal number of clusters is the smallest *k* such that 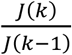 is smaller than certain critical value (Milligan & Cooper, 1985). In our case of binary distances, we limit ourselves to request that ratio to be minimal, indicating a possible correct number of clusters, and define *J*(*k*) as a “sum of mismatching similarity distances error”:

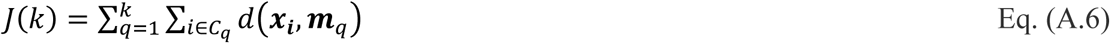

-The *Silhouette* width (Kaufman & Rousseeuw, 1990) is calculated with the following expression:

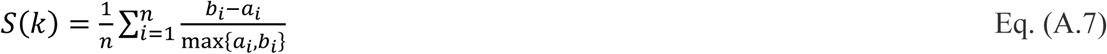

here, *a*_*i*_ is the average distance from the *i*th point to every other object in its cluster: 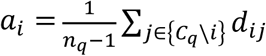 and*b*_*i*_ is the minimum average distance from the *i*th object to all objects of other clusters, minimized over the clusters, namely: 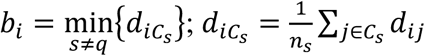(Charrad et al.,2014). The index can take values in the interval [−1,1] with negative values indicating the clustering solution is not accurate, and understandably so, as the minimum average distance from many objects to other clusters would be bigger than the dissimilarity to objects of the clusters where they belong to. On the other hand, the maximum value is taken to represent the optimal number of clusters in the data (Charrad et al., 2014).

-The *Dunn* index (Dunn, 1973) is generally defined as:

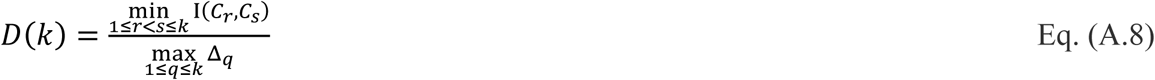

where Δ_*q*_ is the diameter of the *q*th cluster and I(*C*_*r*_, *C*_*s*_) is the intercluster distance between *C*_*r*_ and *C*_*s*_. Out of the many variants available for computing both these quantities, we use the ones recommended by (Bezdek & Pal, 1998):

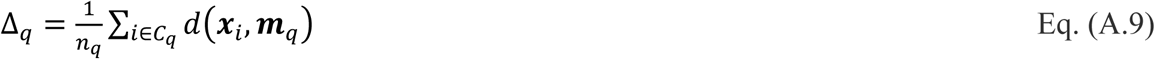

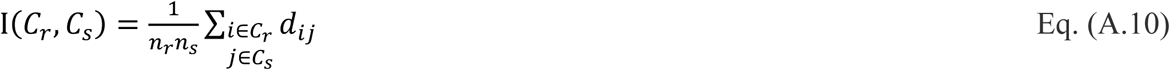

*D* is maximized when the clusters are compact (the diameter is small) and separate (the intercluster distance is large) (Milligan & Cooper, 1985).

-The *Davies-Bouldin* index (Davies & Bouldin, 1979) is also a function of the ratio of within-cluster dispersions and the between-clusters separation. When using mismatching dissimilarities, it can be calculated as:

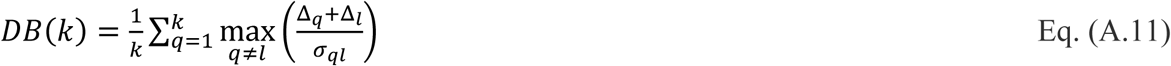

where Δ_*q*_ is defined as in Eq. (A.9) and σ_*ql*_ is the distance between the centroids of clusters *C*_*q*_ and *C*_*l*_, σ_*ql*_ = *d*(***m***_*q*_, ***m***_*l*_) (Charrad et al., 2014). The smaller *DB*(*k*), the better the partition (Dubes, 1987).

The diameter, Δ_*q*_, is zero for clusters with single members. Thus, as highlighted by Davies & Bouldin, theirs and most measures, have limited meaning for singleton clusters.

## Appendix C. Decision rules for random data

The above-mentioned decision rules are not adequate for identifying the correct number of clusters in the limit case of two versus one cluster, as such indexes are not defined for partitions of a lone community (Dubes, 1987). Although the Duda-Hart index was originally designed to reject the existence of only one cluster in the data, the critical value used for such means was obtained by supposing that data came from a normal distribution (Duda et al., 2001), which does not hold in our case of binary variables. Two of the other measures had either simplistic or well-established criteria that could be applied when they and the majority of the indexes indicated the presence of two communities. The first one is inspecting for negative silhouette values (Kaufman & Rousseeuw, 1990). The second rule checks for the presence of a significant drop in the curve of *DB*(*k*) at *k* = 2 (Dubes, 1987). A user can conclude one cluster is present when *DB*(*k*) has a

minimum at 2 but

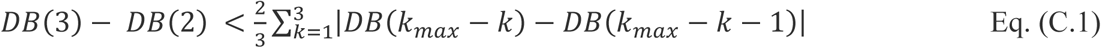

These two criteria were applied to resolve the optimal number of clusters in the limit case situation over all the networks. Consistently, spurious two-clusters partitions over different IMFs were rejected (one single cluster existed). On the contrary, meaningful or real two-communities patterns were indicated as correct by our method and restated by the silhouette and the *DB*-based limit-case criteria.

## Appendix D. Estimating the similarity of partitions

To determine the effectiveness of our clustering technique in retrieving a planted structure, we computed the adjusted mutual information, which establishes a measure of similarity between two partitions based on information theory while adjusting for chance (N. Vinh et al., 2010; Weir, Emmons, Gibson, Taylor, & Mucha, 2017):

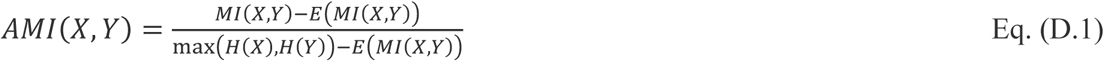

where *MI*(*X, Y*) is the mutual information between random variables *X* and *Y, H*(*X*) is the entropy of *X* and the expected mutual information, *E*(*MI*(*X, Y*)), is obtained as in (N. X. Vinh, Epps, & Bailey, 2009). A MATLAB implementation of *AMI* is available from the Network Community Toolbox (http://commdetect.weebly.com/).

## Acknowledgments

We thank the anonymous referees for the suggestions made during the review process. We are grateful to Anthony Winder for his comments on the manuscript. We also thank the High-Performance Computing group at the University of Calgary (UofC) for their support accessing UofC’s computing resources and the Human Connectome Project for assistance with using their data. Data were provided [in part] by the Human Connectome Project, WU-Minn Consortium (Principal Investigators: David Van Essen and Kamil Ugurbil; 1U54MH091657) funded by the 16 NIH Institutes and Centers that support the NIH Blueprint for Neuroscience Research; and by the McDonnell Center for Systems Neuroscience at Washington University.

## Funding resources

This work was partially supported by the Natural Sciences and Engineering Research Council of Canada (RGPIN-2015-05966).

## Footnotes

1 The terms employed here are mostly faithful to the ones used in each of the specific literature, combined. This is why the word mode appears with two meanings: one refers to the oscillatory functions in which a signal can be decomposed (the intrinsic mode functions), the other represents ‘the centroids’ of the clusters of nodes obtained with binary features (as in *k*-modes clustering).

## Notes

### Competing Interest Statement

The authors have declared no competing interest.

### Summary of Updates

Section "Interpreting the communities identified by the method" added to demonstrate the relationship between network spatial scales and the oscillatory modes of the network signal (Figure 8 added); Figure 7 added to show the application of the algorithm to human brain resting-state fMRI network; Supplemental files updated.

