## Supplementary material for "Detecting brain network communities: considering the role of information flow and its different temporal scales"

### Appendix A. Supplementary material

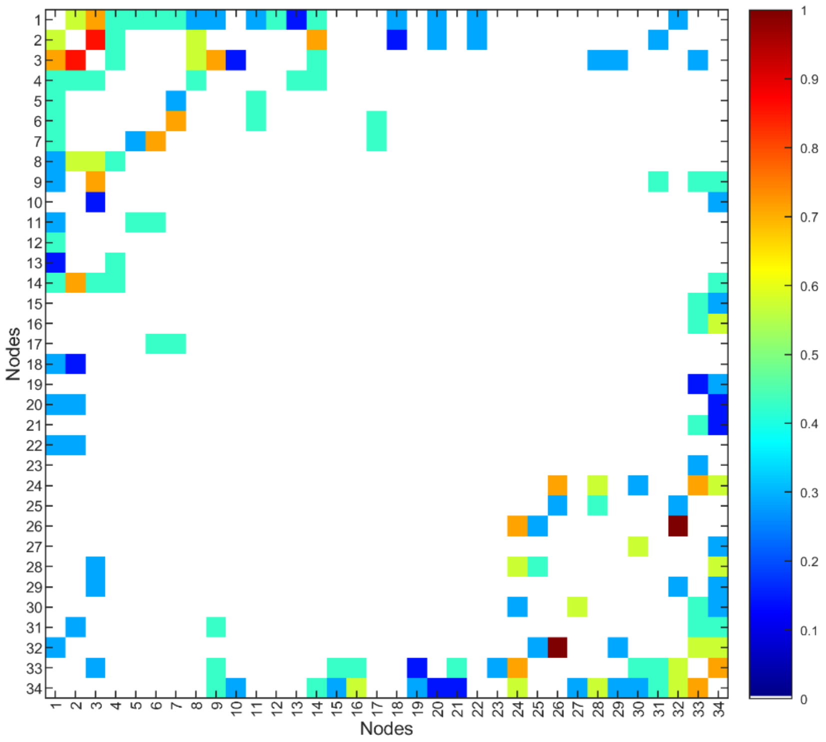


**Fig. S1.** Matrix of interactions in the karate club. The values indicate the number of contexts in which two members of the club were seen interacting, normalized to the interval [0, 1] (dividing by the maximum value, 7). Several connections present inconsistencies in Zachary's original paper. When two values in symmetric positions differed, we chose the lowest one. See also: Zachary, W. W. (1977). An Information Flow Model for Conflict and Fission in Small Groups (https://www.jstor.org/stable/3629752).

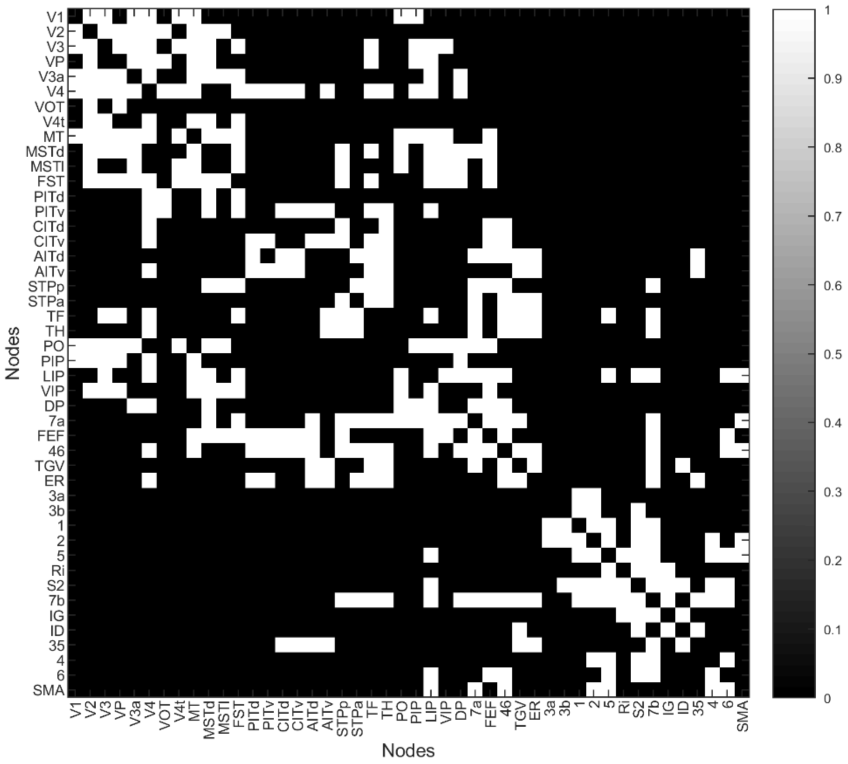


**Fig. S2.** Connectivity matrix of the macaque neocortex. Pathways that have been identified in tracing studies are marked by 1's. See also: Young, M. P. (1993). The organization of neural systems in the primate cerebral cortex (http://doi.org/10.1098/rspb.1993.0040) and Honey, C. J., Kotter, R., Breakspear, M., & Sporns, O. (2007). Network structure of cerebral cortex shapes functional connectivity on multiple time scales (http://doi.org/10.1073/pnas.0701519104).

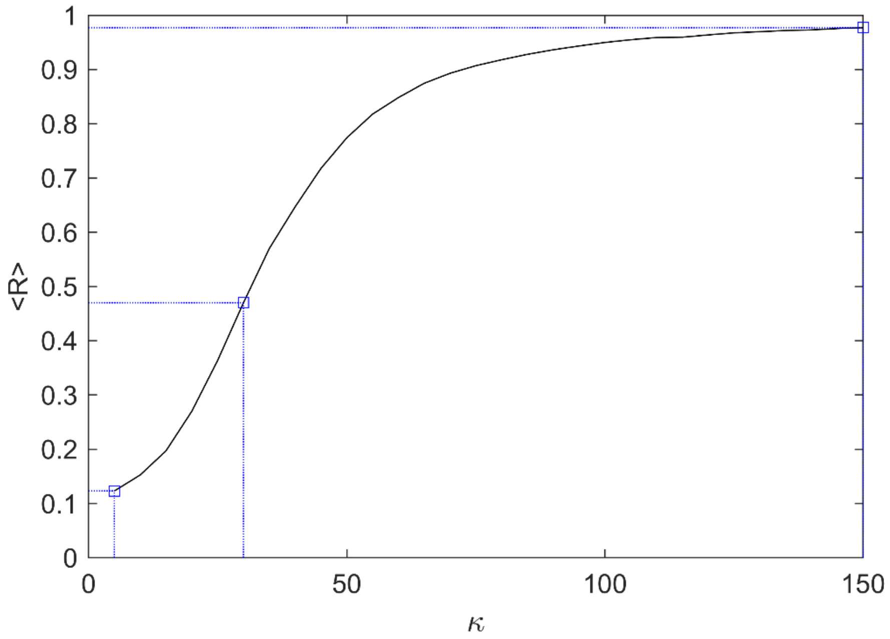


**Fig. S3.** Grand average of the phase uniformity value, $R$ vs the coupling strength, $\kappa = 0{:}\,5{:}\,150$, of the Kuramoto oscillators. The blue squares represent the values of $\kappa$ for which the correlation matrices shown in Fig. 3 were calculated.

| Independent component | Yeo 17-sub-systems | Label |
|---|---|---|
| 1 | 2 | Visual 2 |
| 2 | 16 | DMN 3 |
| 3 | 1 | Visual 1 |
| 4 | 1 | Visual 1 |
| 5 | 13 | Frontoparietal 3 |
| 6 | 1 | Visual 1 |
| 7 | 17 | DMN 4 |
| 8 | 6 | Dorsal attention 2 |
| 9 | 12 | Frontoparietal 2 |
| 10 | 12 | Frontoparietal 2 |
| 11 | 15 | DMN 2 |
| 12 | 7 | Ventral attention 1 |
| 13 | 2 | Visual 2 |
| 14 | 1 | Visual 1 |
| 15 | 5 | Dorsal attention 1 |
| 16 | 3 | Somatomotor 1 |
| 17 | - | - |
| 18 | - | - |
| 19 | 11 | Frontoparietal 1 |
| 20 | 4 | Somatomotor 2 |
| 21 | 8 | Ventral attention 2 |
| 22 | 3 | Somatomotor 1 |
| 23 | - | - |
| 24 | 4 | Somatomotor 2 |
| 25 | 7 | Ventral attention 1 |

**Table S1.** Equivalence between the independent components (nodes in the parcellation of the resting-state fMRI) and established functional sub-systems. The first column represents the nodes and the second column, the specific functional sub-system among the 17 identified by Yeo et al., 2011 (liberal mask). In the third column, we show the label corresponding to each of the Yeo cortical systems according to conventional labeling of 7 coarser functional sub-systems. See also: Sacchet, M. D. et al. (2016). Large-scale hypoconnectivity between resting-state functional networks in unmedicated adolescent major depressive disorder (https://doi.org/10.1038/npp.2016.76).

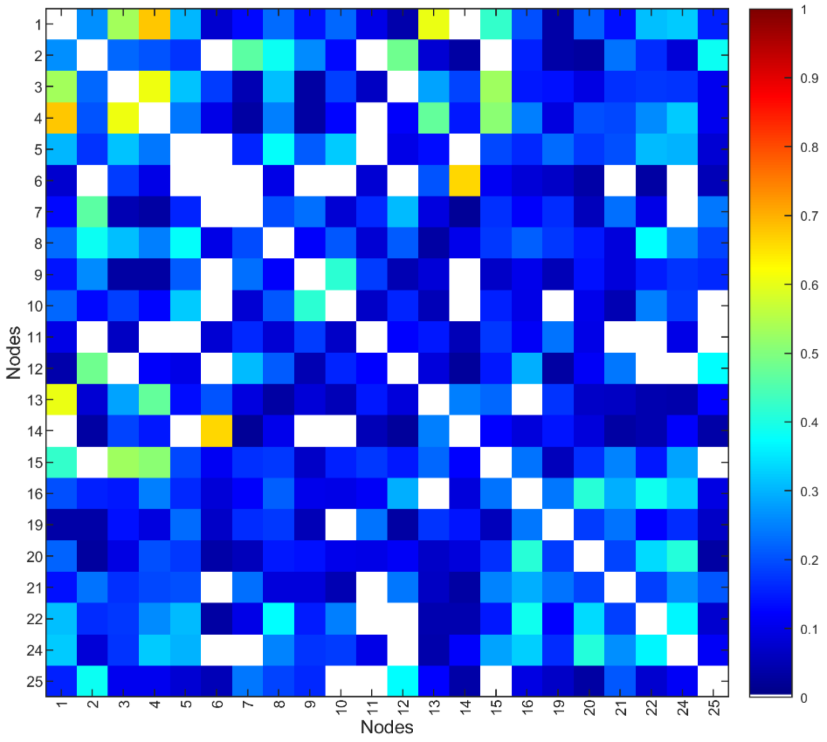


**Fig. S4.** Resting-state functional connectivity matrix in the healthy human brain. Each of the nodes represent a spatial map obtained from performing a group-ICA decomposition of resting-state fMRI. The correlation matrix is averaged across 461 subjects. See also: Smith, S. M. et al. (2015). A positive-negative mode of population covariation links brain connectivity, demographics and behavior (https://doi.org/10.1038/nn.4125) and Smith, S. M. et al. (2013). Resting-state fMRI in the Human Connectome Project (https://doi.org/10.1016/j.neuroimage.2013.05.039).

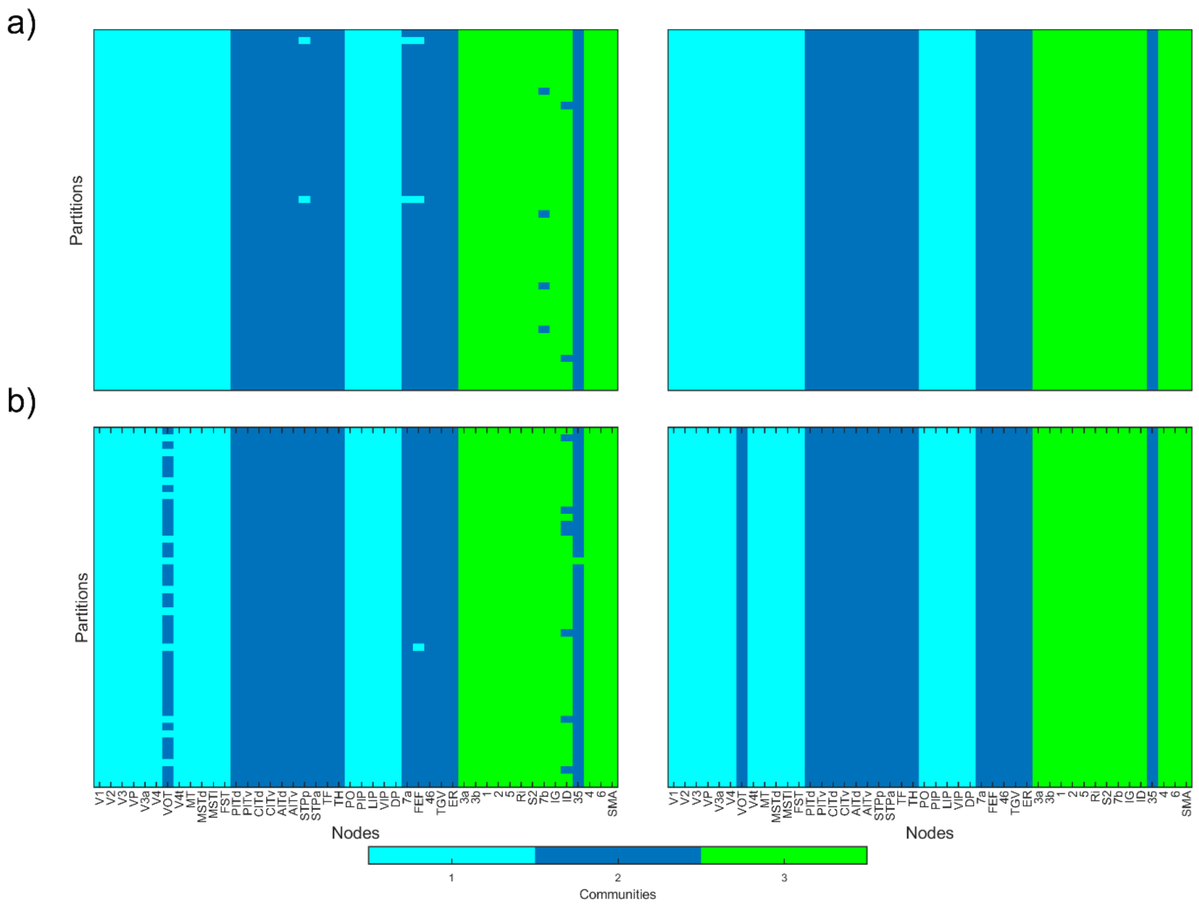


**Fig. S5.** Consensus clustering for the three communities-partitions of the macaque visual and sensorimotor cortex. a) Partitions with the highest similarity (in terms of adjusted mutual information) obtained through runs of the Louvain community detection algorithm (left). This is for $\gamma = 0.9$. Consensus partitions (right). The consensus partition for the more conventional tuning of the resolution parameter, $\gamma = 1.0$, is identical to the one shown. b) Partitions returned by our clustering algorithm with 50 different sets of random features (left) and their consensus clusters (right).

| **Left frontal-cingulate-basal** | **Right frontal-cingulate-basal** | **Left parietal-occipital-temporal** | **Right parietal-occipital-temporal** | **Mixed functional** |
|---|---|---|---|---|
| caudal anterior cingulate | caudal anterior cingulate | cuneus | cuneus | L cuneus |
| caudal middle frontal | caudal middle frontal | entorhinal | entorhinal | L isthmus cingulate |
| lateral orbitofrontal | lateral orbitofrontal | fusiform | fusiform | L lateral occipital |
| medial orbitofrontal | medial orbitofrontal | inferior parietal | inferior parietal | L lingual |
| paracentral | paracentral | inferior temporal | inferior temporal | L pericalcarine |
| pars opercularis | pars opercularis | isthmus cingulate | isthmus cingulate | L precuneus |
| pars orbitalis | pars orbitalis | lateral occipital | lateral occipital | L superior parietal |
| pars triangularis | pars triangularis | lingual | lingual | R cuneus |
| posterior cingulate | posterior cingulate | middle temporal | middle temporal | R isthmus cingulate |
| precentral | precentral | parahippocampal | parahippocampal | R lateral occipital |
| rostral anterior cingulate | rostral anterior cingulate | pericalcarine | pericalcarine | R lingual |
| rostral middle frontal | rostral middle frontal | precuneus | precuneus | R pericalcarine |
| superior frontal | superior frontal | superior parietal | superior parietal | R precuneus |
| insula | insula | superior temporal | superior temporal | R superior parietal |
| accumbens area | accumbens area | supramarginal | supramarginal | R posterior cingulate |
| basal forebrain | basal forebrain | transverse temporal | transverse temporal | |
| caudate | caudate | amygdala | amygdala | |
| putamen | putamen | hippocampus | hippocampus | |
| | postcentral | postcentral | | |
| | pallidum | pallidum | | |
| | thalamus proper | thalamus proper | | |

**Table S2.** List of the brain areas per communities in the organization of the human brain network. The first four columns present the arrangement of four-clusters found in the anatomical network (Fig. 6b). The last column lists all the areas that group inter-hemispherically in the functional network with global coupling strength of the Kuramoto model $\kappa = 5$ (Fig. 6c). The table cells of areas spoiling a completely symmetrical organization are shaded. See also: Klein, A., & Tourville, J. (2012). 101 Labeled Brain Images and a Consistent Human Cortical Labeling Protocol (http://doi.org/10.3389/fnins.2012.00171) and Iturria-Medina, Y., Sotero, R. C., Toussaint, P. J., Mateos-Perez, J. M., Evans, A. C. (2016). Early role of vascular dysregulation on late-onset Alzheimer's disease based on multifactorial data-driven analysis (http://doi.org/10.1038/ncomms11934).

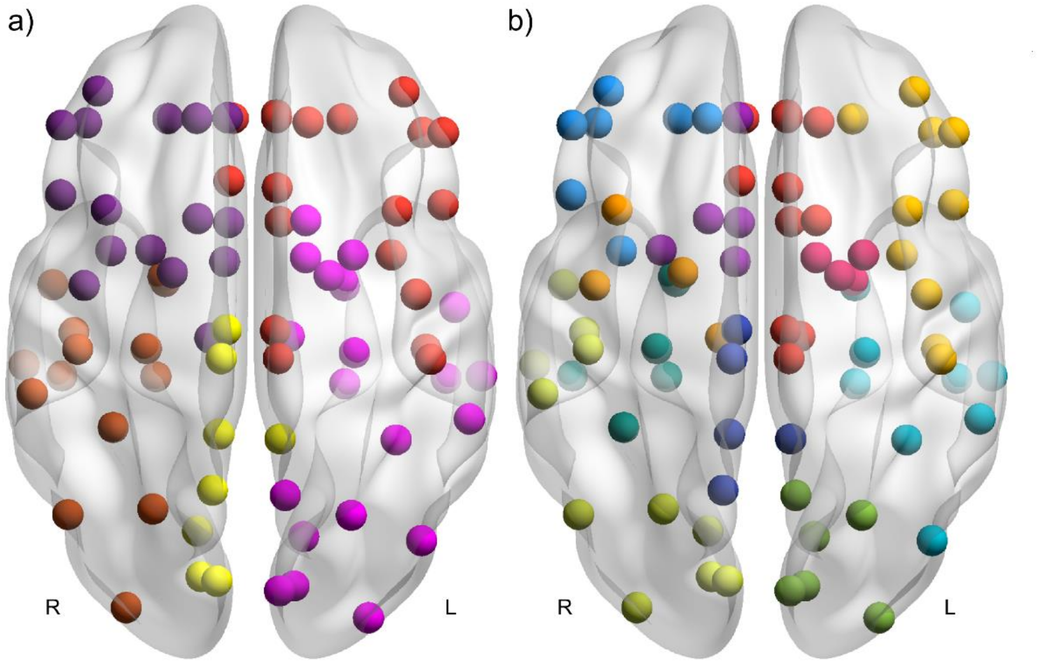


**Fig. S6.** Communities of the human brain functional network ($\kappa = 5$) obtained through Louvain-modularity maximization. a) Partition resulting from taking the resolution parameter $\gamma = 1.0$. b) Partition with the highest similarity (in terms of adjusted mutual information), existing at $\gamma = 3.0$. Colored nodes correspond to communities and their location, to average coordinates of the brain regions in MNI space.

Visualization of the community structures was achieved by means of BrainNet Viewer (http://doi.org/10.1371/journal.pone.0068910).

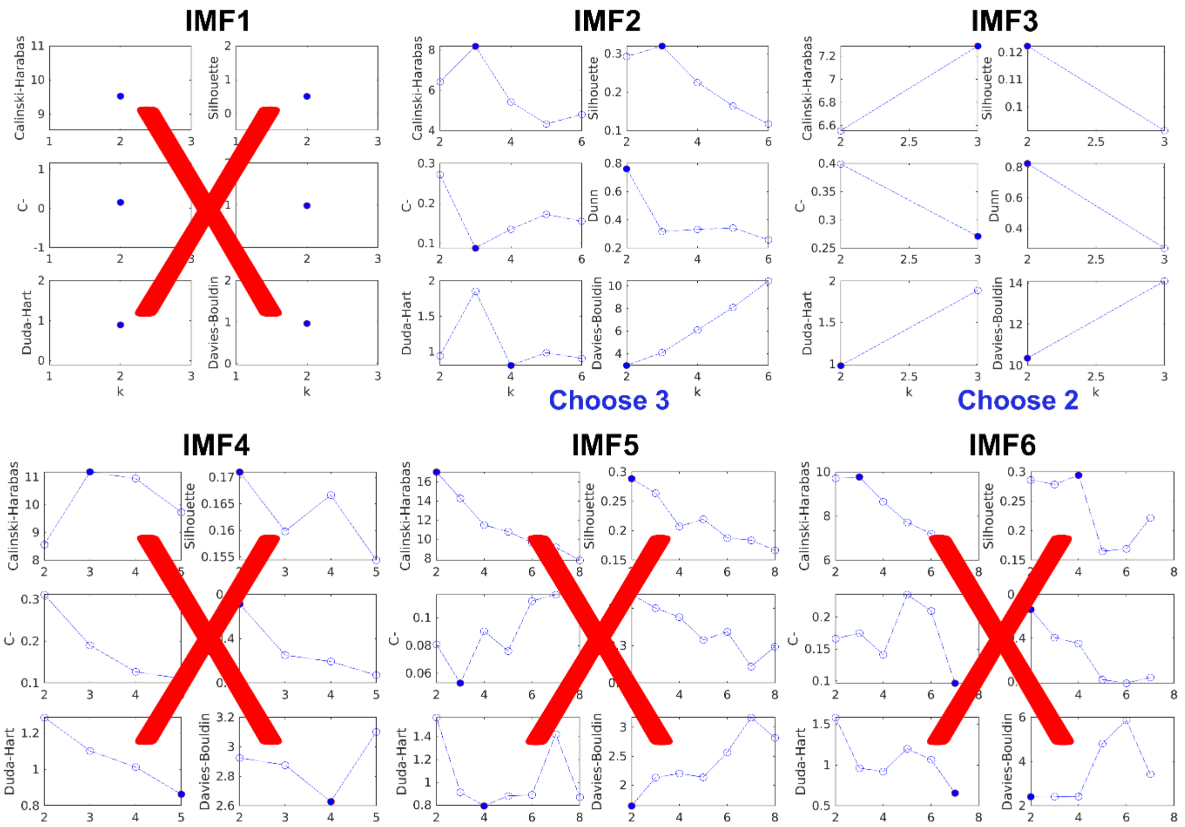


**Fig. S7.** Accepting/rejecting the patterns of communities identified at each IMF. The figure shows the validation indexes of *Appendix B*, computed for the clusters and features of the IMFs in Zachary's karate club network, as signalized at the top of each panel. Red crosses have been used to indicate partitions (IMFs) that are rejected. In the cases of IMF$_6$ and IMF$_4$, these are discarded because no number of clusters achieved a majority of the validation indexes suggesting nodes grouping in such patterns. IMF$_5$ and IMF$_1$ were neglected based on the *Davies-Bouldin* index's decision rule (*Appendix C*). Particularly, IMF$_1$ produces meaningless singleton communities at all possible number of clusters but 2. This is because the oscillations at this mode are fast and most zero-crossings contain only individual nodes (see, for example, Fig. 1). IMF$_2$ and IMF$_3$ do produce meaningful clusters according to the validation indexes, as reported in the main document. See also: *Methods*, *Accepting/rejecting hierarchical partitions* and *Appendices B* and *C*.

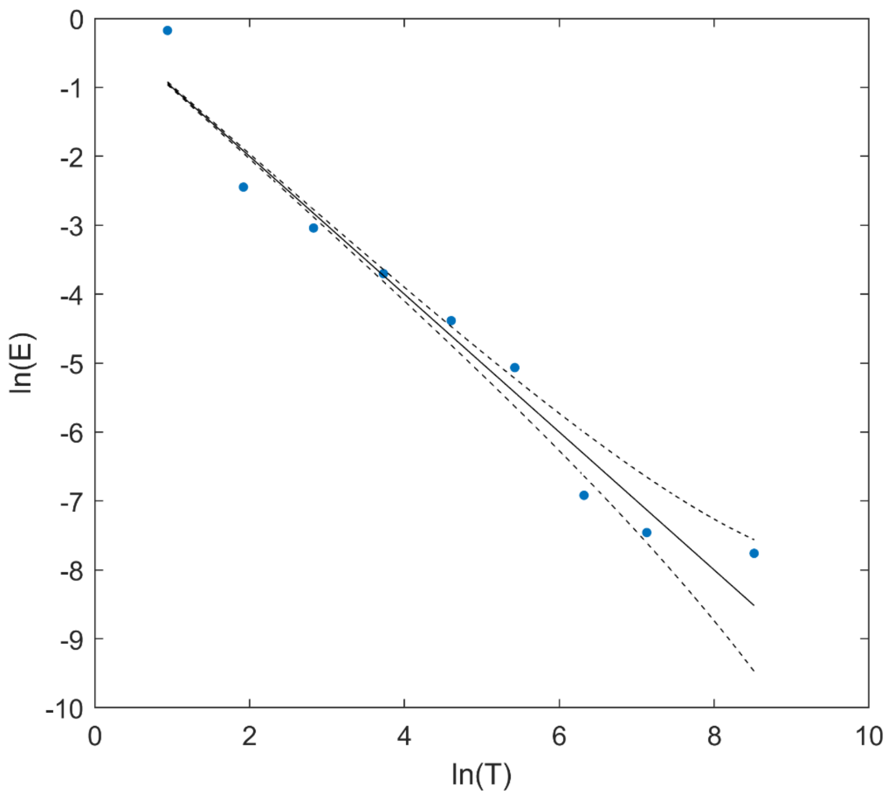


**Fig. S8.** Assessing the information content of the IMFs obtained from decomposing the random walker signals in the karate club network. The figure shows the results of running the statistical test for differentiating IMFs from noise developed by Wu and Huang. The dots represent the test values for each IMF, starting from $IMF_1$ on the left until $IMF_9$ in the right-most part. IMFs having their energy (vertical axis) located above the upper dashed line and below the lower dashed line are considered as containing information at a 99% confidence level. $IMF_4$ was consistently identified as noise across the walkers. However, the meaningful reported partitions appeared over different IMFs. The two-communities ground-truth partition of the karate club network was obtained with the features of $IMF_3$, the next partition being of three communities over $IMF_2$. See also: Wu, Z., & Huang, N. E. (2004). A study of the characteristics of white noise using the empirical mode decomposition method (https://doi.org/10.1098/rspa.2003.1221).

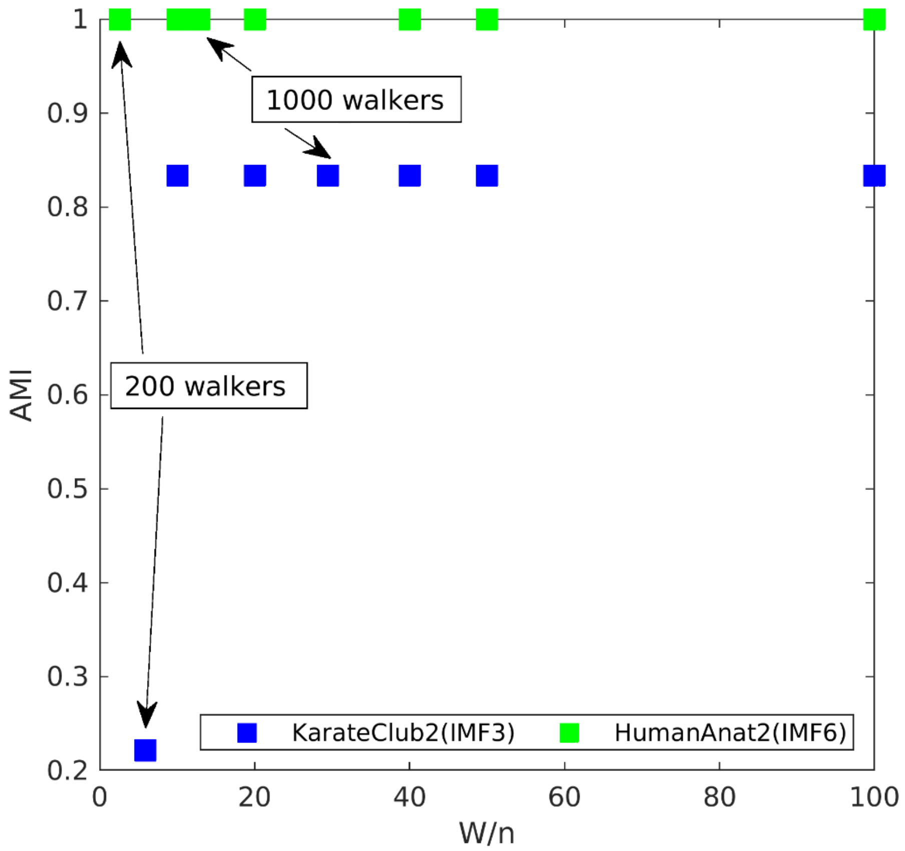


**Fig. S9.** Robustness of the method with regards to the sizes of the networks and number of random walkers used. Values of adjusted mutual information between the returned communities and the ground-truth partitions are plotted for Zachary's karate club (34 nodes; $IMF_3$) and the human brain anatomical network (78 nodes; $IMF_6$). Both curves start from their respective value corresponding to the ratio number of walkers/number of nodes ($W/n$) of $W = 200$, which is the minimum number of walkers employed to compute clustering features. The other point at which the fraction $W/n$ has different values for these two networks is the one corresponding to $W = 1000$ (the adopted standard value in the calculations). The range is thoroughly explored up to $W/n = 50$, while $W/n = 100$ is offered as well. The algorithm did not return Zachary's ground-truth communities (but node 9, See Fig. 4) for $W = 200 < 10 \cdot n$.

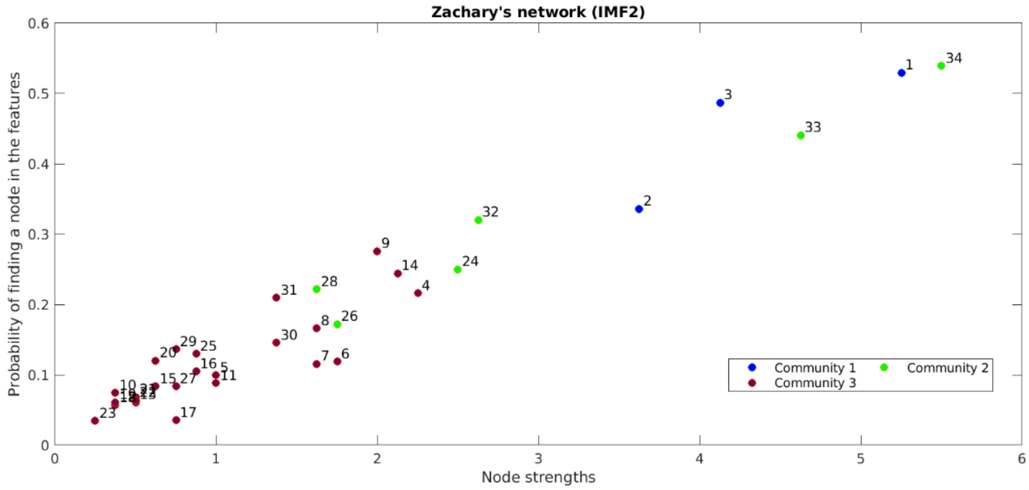


**Fig. S10.** Exploring the influence of the networks' topological properties in the clusters returned by the algorithm. For the heterogeneous Zachary's karate club network, we show the relationship between the probability of finding each node in the clustering features and the total strength of the links the node has, at $IMF_2$. Since nodes 1 and 34 (the two leaders of the club) had more connections than the rest, one may expect that they (and perhaps their close followers) appear more often than the rest when a random walker travels across the network, thus provoking the low-strength nodes to cluster by default. Although some nodes in 'community 3' appear rarely and group together, as shown, nodes 4, 6, 7, 8, 9 and 14 have higher strength than nodes belonging to the leaders' clusters, while 4, 9, 14 and 31 appear more often. This demonstrates that although the node strength contributes to the grouping of nodes, other topological factors also determine communities obtained through our algorithm. If the nodes grouping were based on the strengths only, nodes 26 and 28 would definitely be identified as members of 'community 3'. See also Fig. 4 and its discussion in the main document for an interpretation of the role of each of these communities in the dynamics of the social network.

| Network | No. Nodes | Sparsity | No. Communities |
|---|---|---|---|
| Brain functional | 22 | 0.16 | 5^ |
| Macaque | 46 | 0.79 | 3^ |
| LFR (binary) | 100 | 0.87 | 5*,^ |
| Zachary's | 34 | 0.87 | 2*,^ |
| Brain anatomical | 78 | 0.91 | 4^ |

**Table S3.** Analysis of the number of communities returned by the method vs the characteristics of each network. The networks have been ordered according to their sparsity levels ($1 - \frac{m}{n^2}$; $m$ is the number of non-zero links; $n$ is the number of nodes) from denser to sparser. The number of communities identified in the graphs for which ground-truth communities are known have been marked by a ‘ * ’ (shaded cells). The rest of the networks were explored by using our community detection algorithm. The obtained numbers of clusters in all cases carry the symbol ‘ ^ ’.
