## Supplementary figures and images for "Detecting brain network communities: considering the role of information flow and its different temporal scales"

### Supplementary Video

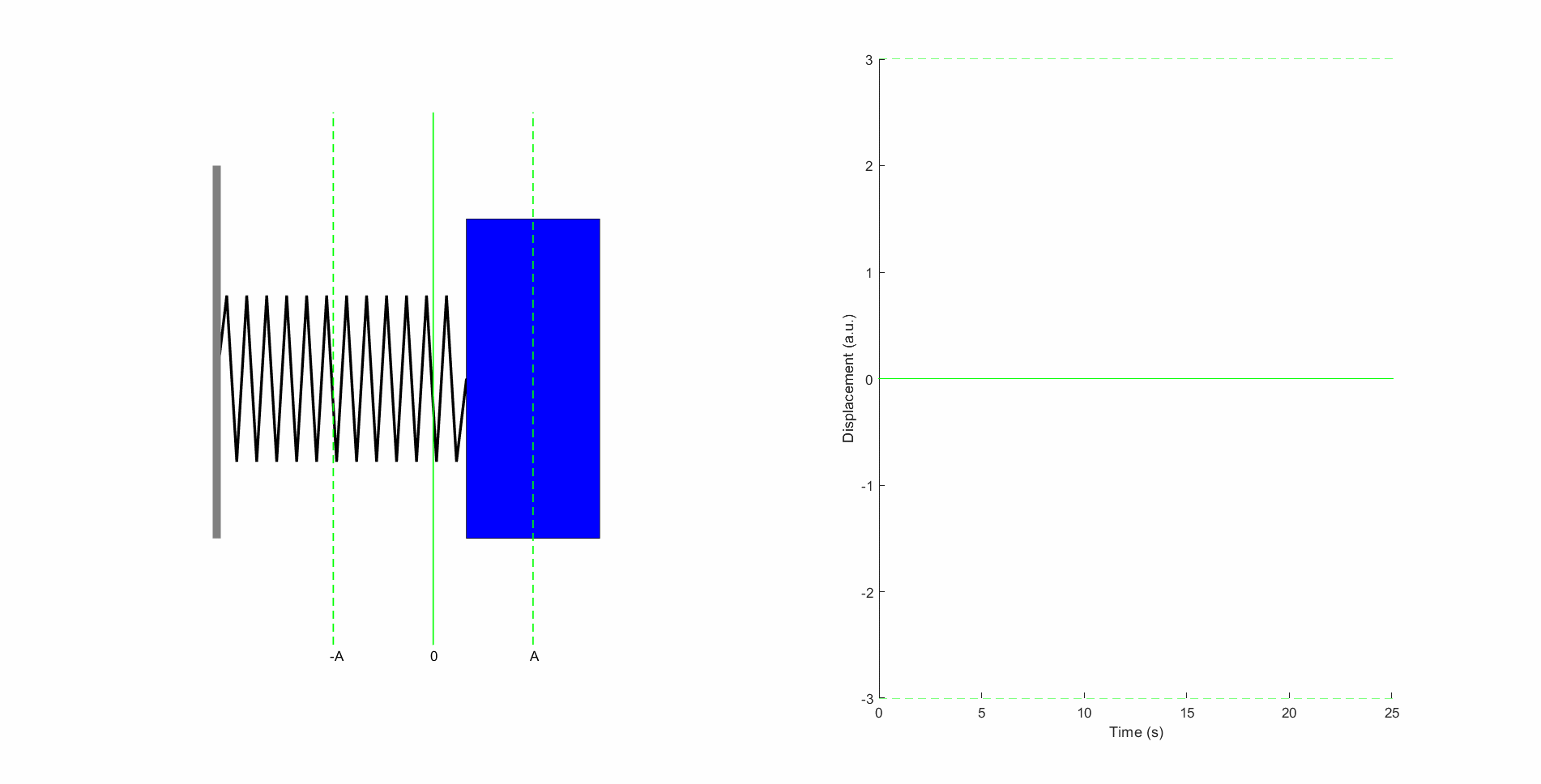
